# HEDeST: An Integrative Approach to Enhance Spatial Transcriptomic Deconvolution with Histology

**DOI:** 10.64898/2026.01.06.697922

**Authors:** Luca Gortana, Loïc Chadoutaud, Raphaël Bourgade, Emmanuel Barillot, Thomas Walter

## Abstract

Spatial organization of cells is essential for tissue function, yet sequencing-based spatial transcriptomics often lacks single-cell resolution. We present HEDeST, a weakly supervised framework that integrates histology-derived morphological features with deconvolution-derived spot-level proportions to assign cell types at single-cell resolution. HEDeST is robust to technical variability, adaptable to user-defined cell types, and compatible with any deconvolution method. Across simulated and semi-simulated datasets, HEDeST outperforms existing morphology-based approaches and reveals biologically meaningful microenvironments when applied to real cancer datasets, providing a scalable tool for high-resolution spatial tissue analysis.

## 1 Introduction

The spatial organization of cells is a defining feature of multicellular life. Cell positioning and interactions maintain tissue homeostasis and drive essential processes such as development [1], immune defense [2], and tissue repair [3]. Pathological conditions often disrupt spatial organization. In cancer, for example, malignant cells reshape the microenvironment by modulating immune infiltration, reorganizing stromal structures, and establishing abnormal cellular niches [4, 5, 6]. Mapping the spatial arrangement of cells is therefore essential for understanding tissue organization and multicellular biology.

Spatial transcriptomics (ST) has emerged as a powerful approach to study cellular organization by measuring gene expression while preserving tissue context [7, 8]. This technology represents a major advance over bulk or dissociated RNA-sequencing (RNA-seq) methods, which lose spatial information. ST methods fall into two broad categories: imaging-based and sequencing-based. Imaging-based technologies rely on fluorescently labeled RNA probes to visualize hundreds to thousands of genes at single-cell or even subcellular resolution [9]. While they provide unmatched spatial resolution, they are limited to predefined gene panels and are less suitable for exploratory, transcriptome-wide analyses. By contrast, sequencing-based ST captures whole-transcriptome information, enabling the study of broad transcriptional programs without prior selection of marker genes [9]. However, their main limitation lies in spatial resolution. In most current implementations, including Visium v1/v2 [7], Slide-seq [10], DBIT-seq [11], and GeoMX DSP [12], each capture area (hereafter referred to as spots) typically encompasses multiple cells. As a result, the measured transcriptome represents an aggregate signal rather than the contribution of individual cells. More recently, technologies such as Visium HD have aimed to increase resolution to the subcellular level [13]. However, in practice, because of high dropout rates, the fine-grained grid often necessitate aggregation across neighboring capture areas, meaning that single-cell resolution is still not systematically achieved.

To overcome the resolution limitation in sequencing-based ST, a variety of computational methods for cell-type deconvolution have been developed [14]. These tools aim to disentangle the mixture of cell types present within each spot, typically by leveraging a single-cell RNA-seq (scRNA-seq) reference [14]. State-of-the-art methods such as Cell2location [15], RCTD [16], DestVI [17], and Stereoscope [18] estimate the proportions or counts of different cell types per spot, providing valuable information about tissue composition. Nevertheless, their outputs remain restricted to spot-level summaries, leaving unresolved the true single-cell spatial arrangement of individual nuclei.

An underexploited opportunity to bridge this gap lies in histology images [14], which are almost always acquired alongside sequencing-based ST. Hematoxylin and eosin (H&E) staining, in particular, contains rich morphological information that reflects cell type, tissue organization, and pathology. Notably, several recent studies have demonstrated that histology alone can predict underlying gene expression profiles [19, 20, 21], revealing that morphological features indeed encode substantial biological information at molecular level. However, while previous studies have made initial attempts to link histology with sequencing-based ST [22, 23, 24, 25, 26, 27], fully systematic integration for single-cell level cell type annotation remains uncommon. Among recent approaches is HistoCell [27], a pretrained model that leverages H&E images to predict cell types at single-cell resolution, with deconvolution-derived ST labels for supervision. However, because ST information is not incorporated at inference, HistoCell relies exclusively on morphological cues. As many cell types and transcriptional states are not distinguishable from histology alone, this inherently limits the precision and scope of its cell-type predictions.

Here, we present HEDeST (Histology-based Enhancement of Deconvolution for Spatial Transcriptomics), a weakly supervised framework that jointly leverages deconvolution-derived ST data and morphological features from histology to predict cell types at single-cell resolution. The method is designed as an add-on to any deconvolution pipeline and can work with either cell-type proportions or counts, making it compatible with existing tools. By training HEDeST on each individual slide, our approach reduces sensitivity to batch or technology-specific effects and remains adaptable across diverse datasets. Besides reaching single-cell resolution within spots, HEDeST also predicts cell types in between spots, filling the gap inherent to the technology. In this study, we introduce the HEDeST framework and evaluate its performance on one simulated, two semi-simulated, and one real ST dataset, all derived from cancer samples. We demonstrate that HEDeST accurately assigns cell types at the single-cell level, outperforms existing morphology-only approaches, and reveals biologically meaningful microenvironments in cancer tissues. By combining deconvolution-derived priors with morphology, HEDeST provides an interpretable, scalable, and easy-to-use solution for advancing the study of tissue organization in health and disease.

## 2 Results

### 2.1 HEDeST framework

We developed HEDeST as a weakly supervised framework to assign cell types to individual cells directly from histology images, guided by deconvolution-derived spot-level proportions (**Fig. 1a**). The method integrates morphological features extracted from high-resolution whole-slide images (WSIs) with information from ST, enabling single-cell resolution annotation without requiring cell-level ground truth.

**Figure 1.**
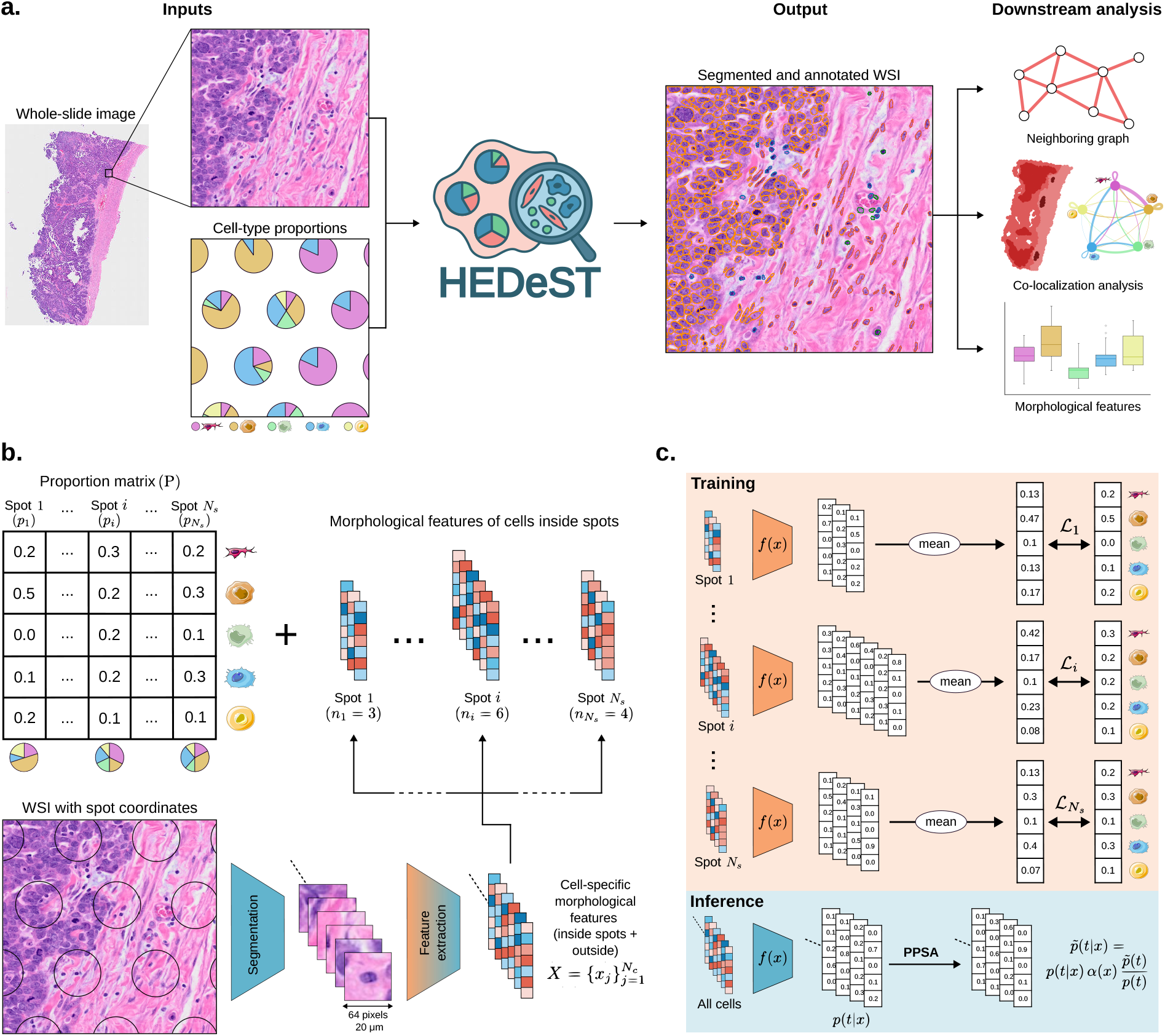
Overview of the HEDeST framework. **a** Schematic overview of the HEDeST workflow. The method integrates histology-derived morphological information with deconvolution results from ST to predict single-cell types across the tissue. HEDeST outputs cell-level annotations and enables downstream analyses such as spatial co-localization, morphological profiling, and neighborhood composition studies. **b** Inputs to the model. Nuclei are first segmented from the H&E image, and cell-centered patches are extracted to compute morphological embeddings. In parallel, spot-level cell-type proportions are obtained from a deconvolution of the ST data. Each cell within a spot is weakly labeled by the corresponding proportion vector. **c** Training and inference strategy. A classifier is trained under a LLP paradigm to align aggregated cell-level predictions with spot-level proportions. After training, the model predicts cell-type probabilities for all nuclei across the slide, including those outside Visium spots. A PPSA step then refines predictions by incorporating local priors. The architecture of the classifier is presented in **Supp. Fig. 1**.

The pipeline relies on two main inputs (**Fig. 1b**). First, WSIs are segmented with HoVerNet [28] to delineate nuclei and extract cell-centered image patches. These patches are then encoded into robust morphological embeddings using self-supervised contrastive learning (MoCo-v3 [29]), such that each cell is represented by a high-dimensional vector encoding/capturing its morphology. Second, spot-level cell-type proportions are obtained from a primary deconvolution of the associated Visium data. HEDeST accepts either proportions or counts (which are internally normalized), ensuring compatibility with any deconvolution tool. Importantly, the set of cell types predicted by HEDeST exactly matches those defined in the primary deconvolution, and thus corresponds to the cell types present in the scRNA-seq reference used for that step (see Methods).

Training proceeds by linking these two inputs under a Learning from Label Proportions (LLP) paradigm [30] (**Fig. 1c**). A cell-level classifier is optimized so that the aggregated predictions within each spot match the known deconvolution-derived composition. The model is trained on each new slide, which minimizes the impact of batch or technology-specific biases. After training, the classifier is applied across the entire slide, yielding a probability distribution over cell types for every nucleus. To further refine predictions, we implemented a Prior Probability Shift Adjustment (PPSA) strategy that incorporates local spot compositions as prior probabilities to calibrate the classifier’s output. For instance, if the spatial transcriptomics data indicates absence of T-cells in a given spot, the PPSA prevents the model from assigning that label, even if the nuclear morphology is suggestive. This mechanism thus ensures that the model resolves one-to-many mappings between morphology and gene expression, leveraging the deconvolution signal to constrain the morphological classifier in ambiguous cases. The method also extends to cells outside Visium spots by interpolating priors from neighboring spots (see Methods).

By providing cell-level predictions, HEDeST allows for tissue characterization at singlecell resolution, analysis of spatial neighborhoods and cell co-localization, and refinement of deconvolution results by recomputing spot-level proportions from cell predictions. Together, these analyses provide a more granular view of tissue organization and microenvironmental interactions than spot-level deconvolution alone.

### 2.2 HEDeST reconstructs morphological cell clusters

We first evaluated the performance of HEDeST in a fully controlled simulation setting, where we generated virtual spots with varying cell type compositions, and ensuring that cell types can be clearly distinguished from the cell embeddings. Using this simplified scenario allows us to assess whether the weakly supervised model is capable of recovering morphological clusters using only spot-level proportions as supervision. In parallel, this step enabled us to determine the best parameters for the model under controlled conditions (see Methods; **Supp. Fig. 2**).

To construct the simulated dataset, we started from the H&E image of the Visium FFPE Human Ovarian Cancer sample [31]. Following nucleus segmentation, morphological embeddings were extracted and subsequently grouped into clusters using the K-means algorithm. To ensure well-separated morphological groups, we retained the 2,000 cells nearest to each cluster centroid. These cells were then randomly aggregated into pseudo-spots, for which the cluster composition was therefore known (see Methods). We considered two conditions: (i) a balanced scenario, in which all clusters were equally represented across the spots, and (ii) a more realistic scenario, characterized by strong imbalance in cluster frequencies (**Fig. 2a**).

**Figure 2.**
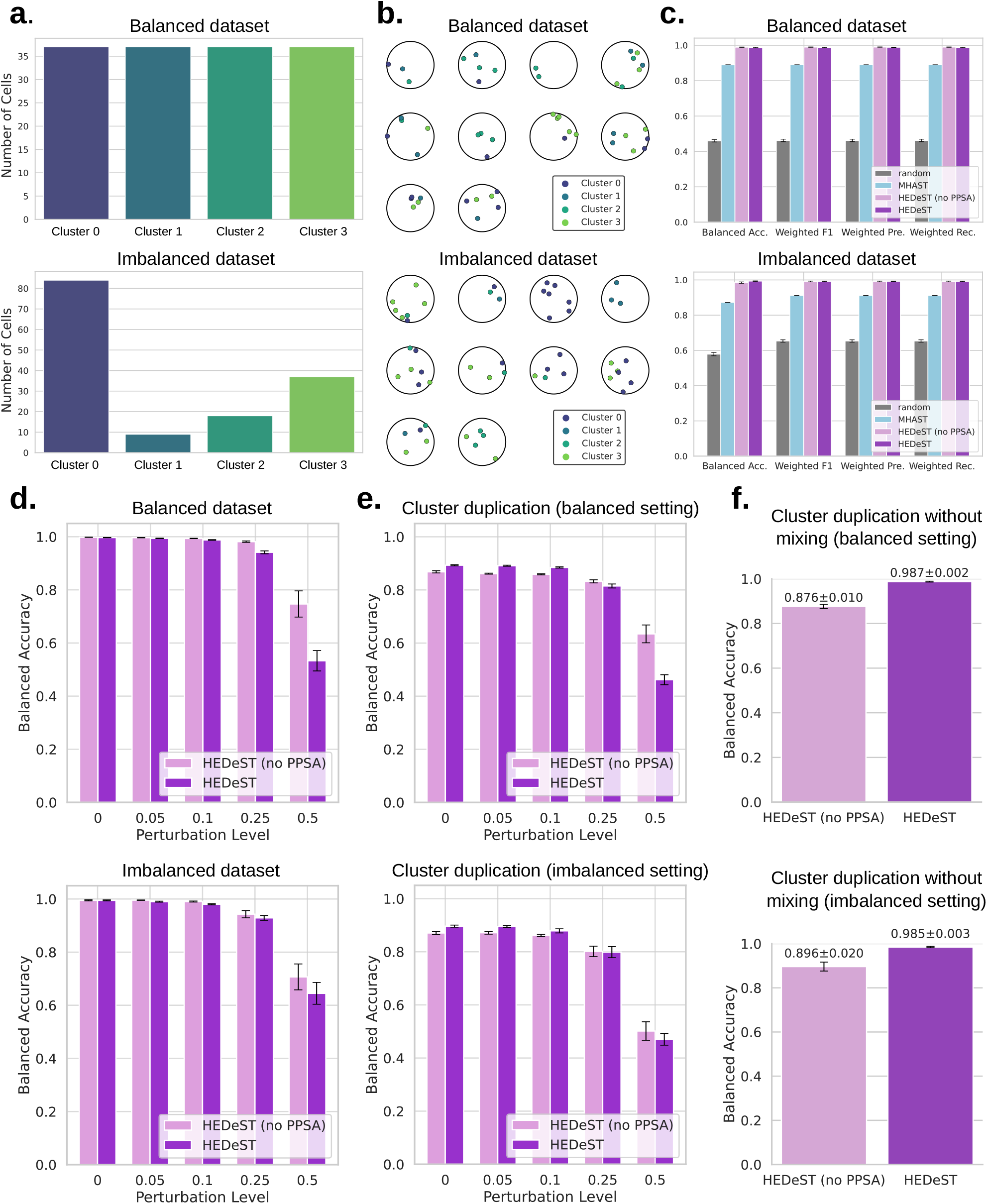
Performance of HEDeST in fully simulated datasets. HEDeST (no PPSA), HEDeST without PPSA; Balanced Acc., Balanced Accuracy; Weighted Pre., Weighted Precision; Weighted Rec., Weighted Recall. **a** Composition of the balanced and imbalanced simulated datasets, each containing four morphological clusters distributed across 30 pseudo-spots. **b** Visualization of the simulated pseudo-spots. **c** Benchmark comparison between HEDeST and MHAST on the balanced and imbalanced scenarios. HEDeST consistently outperformed MHAST in both settings. Bar heights represent the mean across 100 runs, and the error bars indicate 95% confidence intervals computed from the standard error. d Performance of HEDeST on the extended balanced (**top**) and imbalanced (**bottom**) datasets (200 pseudo-spots, 6 clusters) across increasing perturbation levels of pseudo-spot proportions. HEDeST maintains robust performance even under high proportion perturbations. **e** Evaluation under identical conditions as in d, with a duplicated seventh cluster (morphologically identical to cluster 0), demonstrating that PPSA improves accuracy when handling morphologically similar clusters. **f** Comparison of HEDeST with and without PPSA on the balanced (**top**) and imbalanced (**bottom**) datasets where duplicated clusters never co-occur within the same pseudo-spot. No perturbation experiment has been performed here. HEDeST achieves nearly perfect accuracy, indicating that morphologically identical cells can be distinguished via proportion-based context when spatially segregated. Bar heights represent the mean across 10 runs, and the error bars indicate 95% confidence intervals computed from the standard error.

We benchmarked HEDeST against MHAST [32], a method that formulates the problem as an optimization task matching deconvolved spot counts to nuclei through hierarchical permutation. While theoretically well-suited to the task, MHAST is computationally prohibitive due to the exponential scaling of the optimization space, which restricted the experiment to four morphological clusters distributed across 30 pseudo-spots (far from the 4000 spots typically found in a Visium slide), with the number of cells per spot drawn from a normal distribution with mean and variance both equal to 5 (see Methods; **Fig. 2b**). In this setting, HEDeST successfully reconstructed the original morphological clusters from spot-level proportions and consistently outperformed MHAST (**Fig. 2c**). The model achieved a balanced accuracy above 0.98 in the balanced scenario and exceeded 0.99 in the imbalanced scenario. In the balanced case, HEDeST maintained high performance with or without refinement. However, the benefit of the PPSA became particularly evident in the more complex imbalanced scenario, where it improved the balanced accuracy from 0.985 to 0.992.

We next scaled the simulation to 200 pseudo-spots comprising six clusters (mean 15 cells per spot). In this setting, HEDeST again accurately reconstructed the underlying morphological clusters using only spot-level proportions as supervision (**Supp. Fig. 3**). We then introduced increasing levels of proportion perturbation to mimic deconvolution error (see Methods; **Fig. 2d**). Across both balanced and imbalanced datasets, balanced accuracy remained high from low to moderate perturbation, indicating that HEDeST is robust to realistic levels of noise in the priors. As perturbation increased further, performance declined progressively. In the absence of morphological ambiguity, the unadjusted model was more resilient under extreme perturbation, consistent with the fact that PPSA directly incorporates spot-level priors and therefore becomes increasingly penalized when those priors are strongly corrupted. Accordingly, PPSA is most beneficial when deconvolution quality is good or moderately noisy.

To evaluate the role of PPSA under morphological ambiguity, we duplicated one cluster to create an indistinguishable “twin” class (see Methods). In this setting, PPSA provided a clear advantage at low to moderate perturbation levels, outperforming the unadjusted model by leveraging spatial priors to resolve morphologically similar populations (**Fig. 2e, Supp. Fig. 4**). Thus, PPSA is particularly valuable when morphology alone is insufficient and deconvolution estimates remain reasonably reliable. When the duplicated clusters were spatially segregated and never co-occurred within the same spot (**Fig. 2f**), PPSA achieved near-perfect disambiguation. This result highlights that when morphologically indistinguishable clusters are not mixed within the same spatial context, local proportion priors provide enough discriminative information for PPSA to resolve them nearly without error.

Interestingly, under high perturbation, the gap between PPSA-adjusted and unadjusted models was more pronounced in the balanced setting than in the imbalanced one (**Figs. 2d, 2e**). In balanced datasets, all cell types contribute similarly to the spot-level priors; consequently, strong perturbation can considerably distort the compositional signal, generating priors that differ significantly from the true proportions and thereby amplifying differences between HEDeST with and without PPSA. In contrast, in imbalanced datasets, a small number of dominant cell types largely define the priors. Even under extreme perturbation, these majority classes tend to remain predominant in the compositional signal, preserving the overall structure of the dataset and thus explaining the relatively good performance of HEDeST with PPSA. This suggests that PPSA is particularly relevant in realistic biological contexts, such as tumors, where strong cell-type imbalance is common.

Together, these analyses confirm that HEDeST successfully reconstructs the underlying morphological clusters and consistently outperforms a strong baseline method designed for this task. Balanced accuracy remains robust up to moderate perturbation levels, demonstrating resilience to realistic deconvolution noise. The results further highlight that HEDeST, particularly when combined with PPSA, performs best when the deconvolution priors are reliable, and that PPSA is especially valuable for distinguishing cell types with highly similar morphologies.

### 2.3 HEDeST accurately assigns cell types on semi-simulated data

To further assess HEDeST under conditions closer to real data, we constructed semi-simulated datasets from Xenium breast [33, 34] and lung cancer slides [35] (see Methods; **Supp. Figs. 5-15**). In both cases, cell-type labels derived from Xenium data (annotations provided or obtained by scRNA-seq atlas transfer [36]) were mapped onto nuclei segmented on the corresponding H&E images, and pseudo-spots were generated by overlaying Visium-like grids. This produced datasets where the composition of each pseudo-spot was known, enabling quantitative evaluation of HEDeST. Each dataset contained 7 cell types, but with highly imbalanced distributions (**Supp. Figs. 7, 13**), dominated by malignant cells, reflecting the distribution typically observed in tumor samples. Importantly, this semi-simulation step also allowed us to refine the model configuration by selecting the most suitable parameters under realistic conditions (see Methods; **Supp. Fig. 16**).

When trained on these data, HEDeST successfully assigned cell types from morphological features and pseudo-spot-level proportions (**Fig. 3a**). To quantify performance at the spot level, we averaged predicted cell-type probability vectors across all cells within each pseudo-spot. At this resolution, PPSA improved the Pearson correlation (PCC) between predicted and true spot compositions (**Supp. Fig. 17**), illustrating its effect of adjusting predicted probabilities to better match the original spot proportions. At the cell level, galleries of representative single-cell images confirmed that the model learned distinctive morphological signatures for the different classes (**Figs. 3b**). Notably, many cells from the Breast cancer dataset predicted as B cells correspond biologically to plasma cells, which are terminally differentiated B cells.

**Figure 3.**
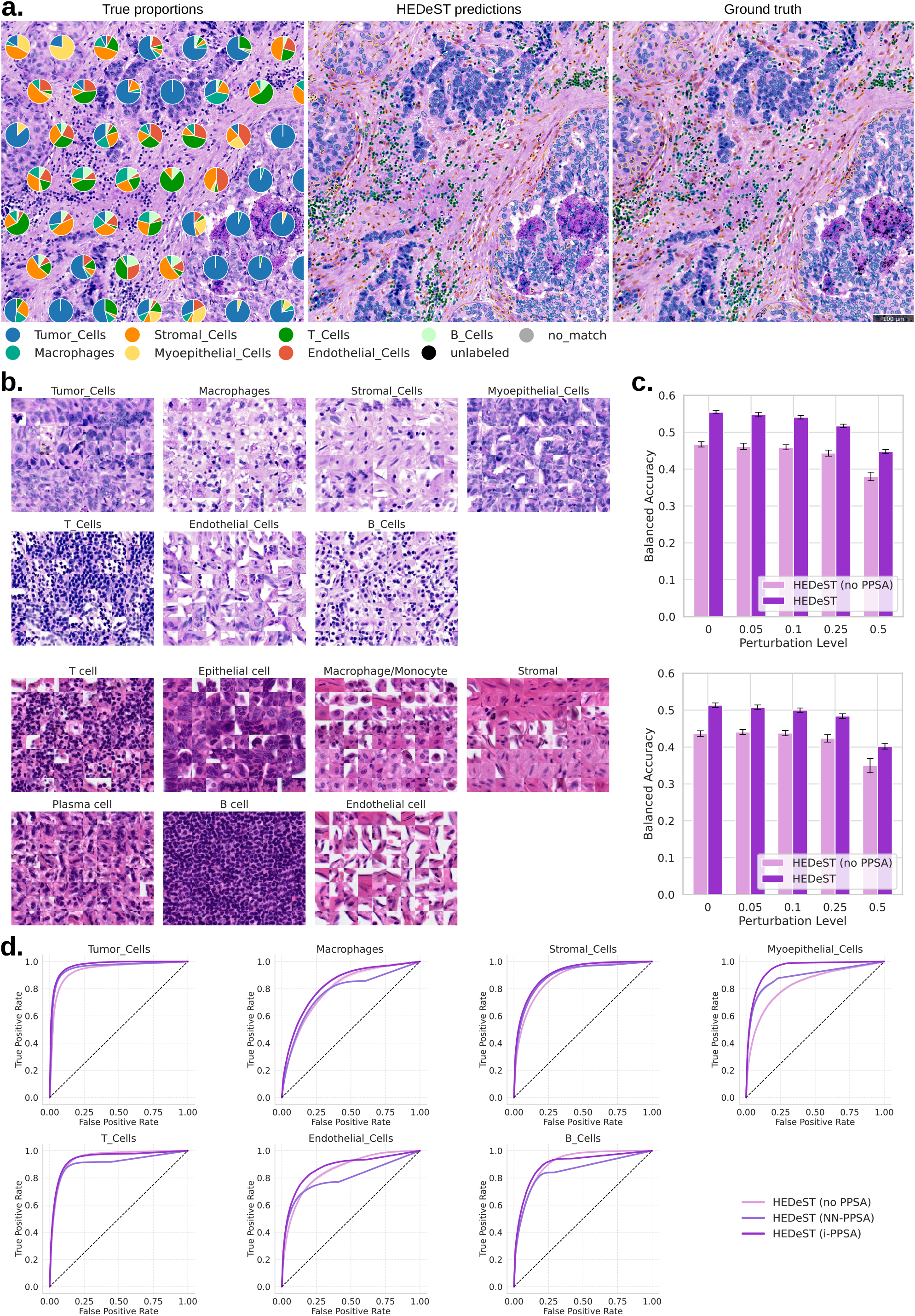
Performance of HEDeST in semi-simulated datasets. HEDeST (no PPSA), HEDeST without PPSA. **a** Visualization of HEDeST predictions on Breast Cancer H&E slide. **Left** H&E image with ground-truth proportions; **Center** segmented and annotated cells based on HEDeST predictions; **Right** segmented and annotated cells based on Xenium ground truth. Scale bar indicates 100 µm. **b** Grid of representative cell crops for each predicted cell type in the Breast (**top**) and Lung (**bottom**) datasets, illustrating consistent and distinct morphological features across all classes. **c** Evaluation of HEDeST performance on the Breast (**top**) and Lung (**bottom**) dataset across increasing perturbation levels of pseudo-spot proportions. Across both tissue types, HEDeST shows robust performance, with PPSA consistently providing the necessary refinement to improve the predictions. Bar heights represent the mean across 5 runs, and the error bars indicate 95% confidence intervals computed from the standard error. **d** Cell-type–specific ROC curves for the Breast Cancer dataset. i-PPSA generated smoother, more biologically consistent curves, avoiding abrupt shifts seen with NN-PPSA.

We repeated the perturbation experiments on the semi-simulated datasets (**Fig. 3c**). Across both breast and lung datasets, PPSA improved performance at all perturbation levels, confirming that PPSA becomes particularly relevant in realistic biological contexts characterized by strong cell-type imbalance and overlapping morphologies. In the absence of perturbation, HEDeST achieved balanced accuracies of 0.554 on the breast cancer slide and 0.513 on the lung cancer slide. Performance was generally higher for cells located inside pseudo-spots, which were well represented in the training set, but HEDeST still achieved reliable predictions for cells outside spots (**Supp. Fig. 18**).

The semi-simulated setting provided a useful testbed to compare alternative strategies for incorporating PPSA within HEDeST. We therefore evaluated three variants: HEDeST without PPSA, HEDeST with PPSA using the nearest-spot proportions as priors for inter-spot cells (NN-PPSA), and HEDeST with interpolated PPSA (i-PPSA), in which priors are defined as a linear combination of the three surrounding spots. ROC curves showed that i-PPSA yielded smoother and more biologically coherent profiles, whereas NN-PPSA introduced abrupt probability shifts that resulted in artificial discontinuities (**Fig. 3d**). The unadjusted model also underperformed the i-PPSA-corrected version. Precision–Recall (PR) curves, which provide a more informative measure under class imbalance, further confirmed the advantages of i-PPSA, particularly for minority cell types where maintaining high recall is crucial (**Supp. Fig. 19**). Based on these results, i-PPSA was adopted as the default PPSA formulation in HEDeST; unless stated otherwise, references to HEDeST correspond to this interpolated variant.

Overall, on semi-simulated datasets, HEDeST successfully combines morphological features with spot-level proportions to achieve accurate single-cell assignments, with PPSA further improving performance at both spot and cell levels. However, the results should be interpreted with caution, as the ground truth itself is imperfect: cell annotations are transferred from Xenium data and single-cell references, both prone to errors. This inaccuracy both constrains the information available to HEDeST during training and can distort the evaluation of predictive performance.

### 2.4 HEDeST outperforms existing cell-type assignment methods

To further evaluate the performance of HEDeST, we benchmarked it against two widely recognized approaches for cell-type inference from histology: HistoCell [27] and HoVerNet [28]. These methods were selected because they reflect two complementary strategies. HistoCell is a deep-learning framework originally proposed as a collection of pretrained models, each specific to one of nine cancer types. The original training relied on weak supervision derived from deconvolution-based cell-type proportions. As the pretrained weights are not publicly available, we retrained HistoCell using our semi-simulated datasets, ensuring that comparisons with HEDeST were made under equivalent conditions (see Methods). This allowed us to directly assess differences in architecture and to evaluate the added contribution of PPSA. HoVerNet is a state-of-the-art method for nuclei segmentation and annotation from H&E images. Trained on curated histology datasets [37, 38], it outputs broad cell-type categories, such as neoplastic, inflammatory, and connective cells. To enable a fair comparison, we adapted our semi-simulated datasets by mapping cell types into the same broad categories and applied HEDeST to this simplified version (see Methods). This approach allowed a conceptual benchmark between pretrained models and slide-specific training strategies.

At the spot level, HEDeST without PPSA and HistoCell demonstrated comparable performances in reconstructing spot-specific cell-type proportions (**Fig. 4a**), and HEDeST with i-PPSA naturally performed much better. At the single-cell level, HEDeST consistently outperformed HistoCell. Balanced accuracy across all evaluated datasets was higher with HEDeST (**Figs. 4b, 4c**). While the overall spot-level predictions were similar, HEDeST produced more reliable distributions of cell-type probabilities across individual cells (**Supp. Fig. 20**), indicating that HEDeST’s architecture and i-PPSA substantially enhanced single-cell classification. In comparison with HoVerNet, HEDeST also achieved superior performance in the 3-class benchmark (**Fig. 4c**), highlighting the advantages of incorporating prior information and retraining on each slide rather than relying on fixed pretrained models. Visual inspection of representative tissue regions confirmed that HEDeST predictions closely matched the ground truth, with more accurate cell-type classification than HistoCell and HoVerNet (**Figs. 4d, 4e**).

**Figure 4.**
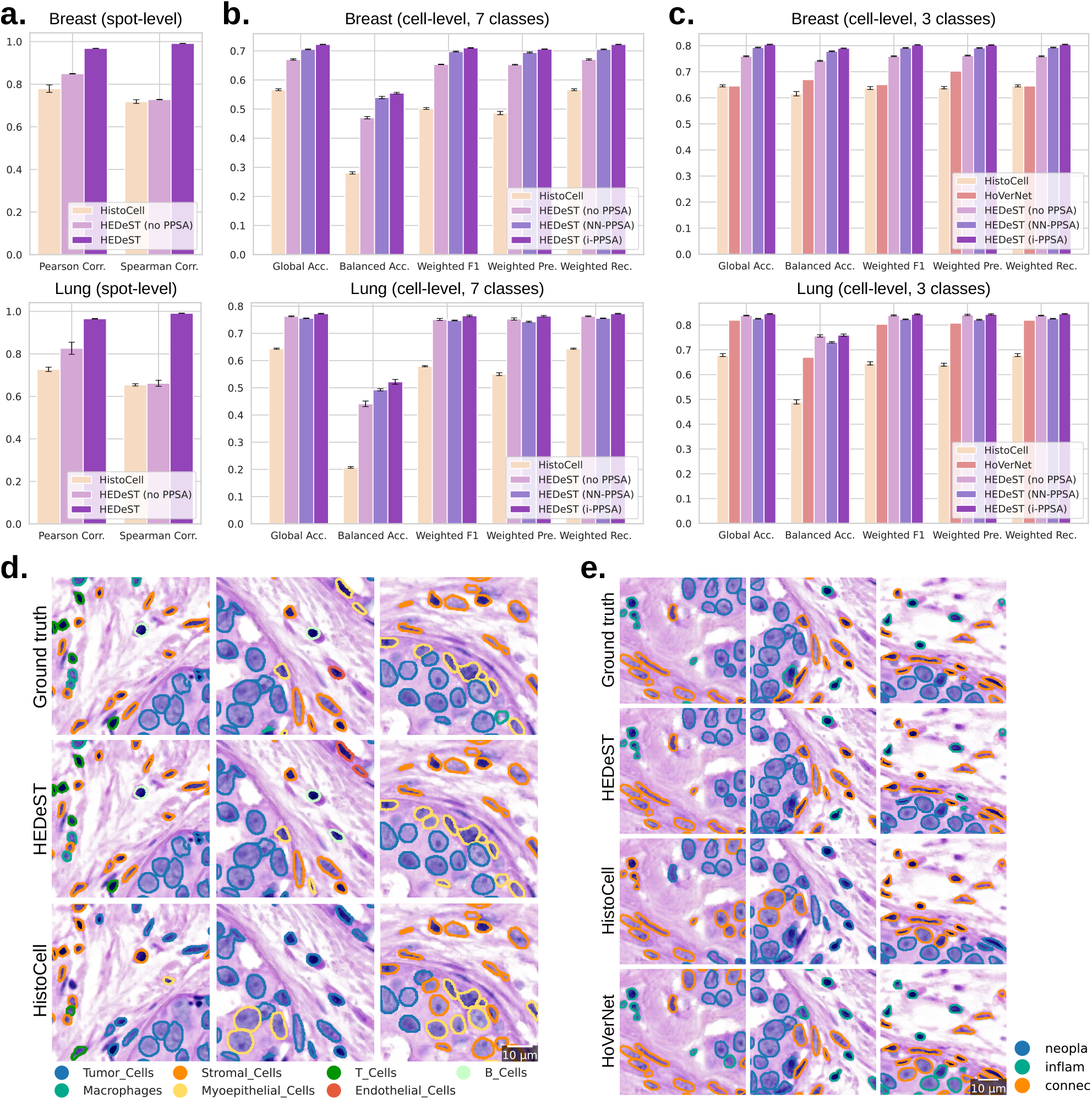
Benchmark performance of HEDeST against state-of-the-art methods. HEDeST (no PPSA), HEDeST without PPSA; Global Acc., Global Accuracy; Balanced Acc., Balanced Accuracy; Weighted Pre., Weighted Precision; Weighted Rec., Weighted Recall; neopla, Neoplastic cells; inflam, Inflammatory cells; connect, Connective cells. **a** Spot-level performance on Breast Cancer (**top**) and Lung Cancer (**bottom**) slides. Global Pearson correlation (aggregated across cell-type–specific correlations between ground truth and predictions) is shown for HistoCell, HEDeST before PPSA, and HEDeST after PPSA (i-PPSA). While HistoCell and HEDeST achieve similar results after training, PPSA further improves HEDeST by better aligning predictions with local spot proportions. **b,c** Cell-level performance metrics computed for datasets with the full set of cell types (b) and restricted to three broad categories (**c**) for both Breast Cancer (**top**) and Lung Cancer (**bottom**) slides. Restricting to three types enabled direct comparison with HoVerNet (see Methods). HEDeST consistently outperformed both HoVerNet and HistoCell across all configurations. Bar heights represent the mean across 5 runs, and the error bars indicate 95% confidence intervals computed from the standard error. **d,e** Representative cell crops comparing HEDeST predictions on breast slide with those from HistoCell, HoVer-Net, and ground truth. Some cells are not outlined or annotated as they were either excluded from semi-simulated datasets or omitted for benchmarking purposes (see Methods). **d** Original semi-simulated datasets; **e** datasets restricted to three cell types. Scale bars indicate 10 µm. HEDeST predictions more closely match ground truth annotations and accurately capture the spatial organization of cells.

In summary, the benchmarking analyses indicate that pairing slide-specific training with PPSA enables HEDeST to outperform pre-trained and image-only methods, making it a more robust and reliable approach for annotating single cell types.

### 2.5 HEDeST reveals single-cell neighborhood organization in breast cancer

To illustrate potential application cases enabled by our framework, we applied HEDeST to a real Visium FFPE breast cancer sample [39] (**Supp. Fig. 21**) obtained from a 73-year-old female patient diagnosed with Grade II Invasive Carcinoma and Ductal Carcinoma In Situ (DCIS). The goal was to investigate whether our framework could provide finer-grained insights beyond deconvolution. Specifically, we sought to assess whether HEDeST could identify cell populations, capture co-localization patterns, and delineate microenvironmental domains that would not be accessible from spot-level data alone.

We performed deconvolution with DestVI [17] using a breast cancer single-cell reference atlas [40, 41] (see Methods), yielding spot-level proportions after post-processing (**Fig. 5a**). HEDeST was then applied to the same slide, producing single-cell type maps (**Fig. 5b, Supp. Fig. 22**). For each predicted class, we inspected representative single-cell images and found that the morphologies corresponded well to known features of the predicted cell types (**Fig. 5c**). To investigate whether HEDeST can be used to reveal morphological characteristics of cell types, we compared nuclear areas across all cell types (**Fig. 5d**). While myeloid cells and plasmocytes generally exhibit larger nuclei, B and T lymphocytes are morphologically similar and difficult to distinguish [42]. Despite these difficulties, we found that nuclei predicted as B cells were significantly larger than those predicted as T cells, in agreement with prior reports [43]. This shows that HEDeST can be used as a tool to identify morphological markers even in cases where phenotypic differences are subtle.

**Figure 5.**
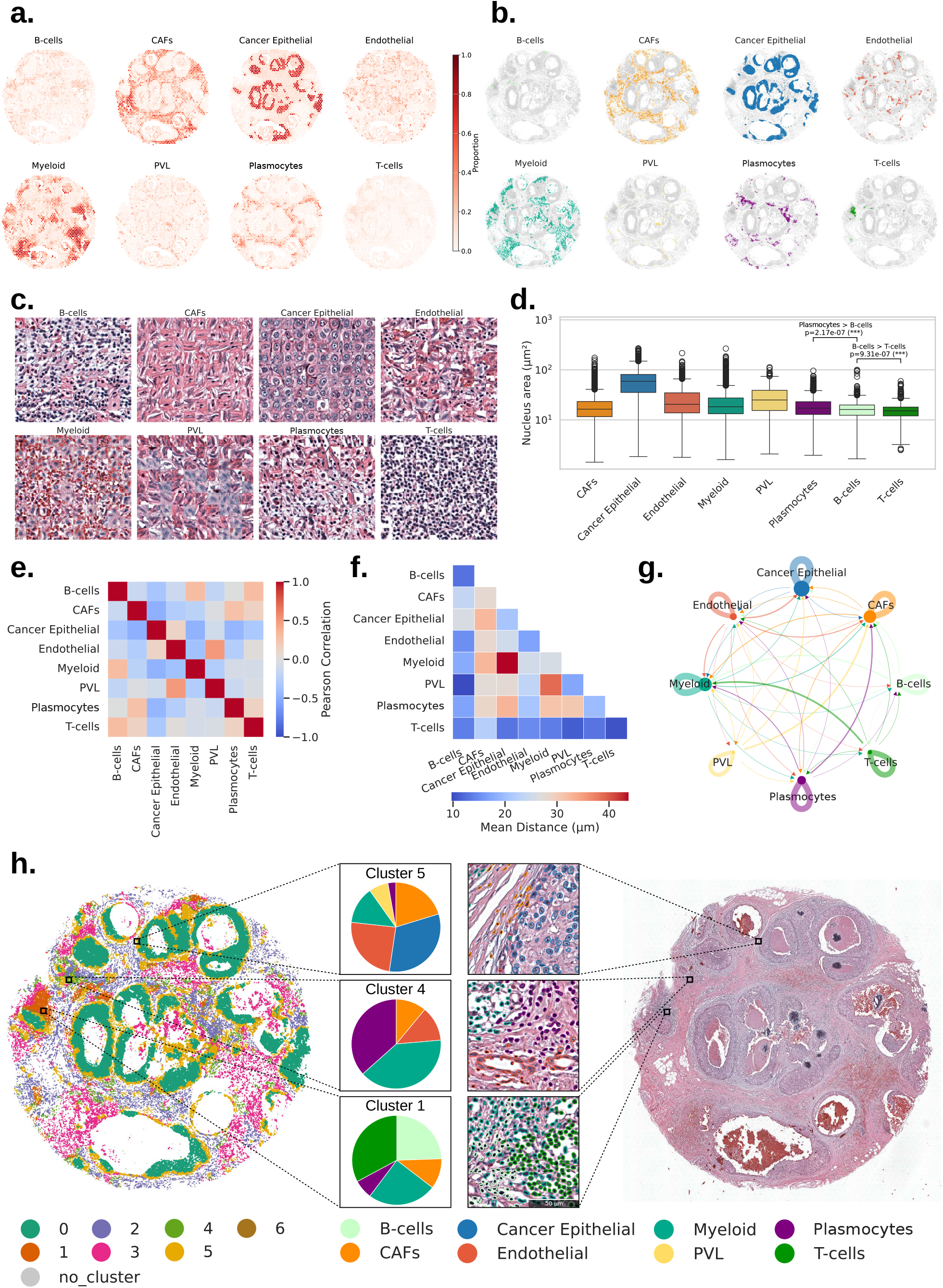
Spatial organization analysis of a real breast cancer sample using HEDeST. CAFs, Cancer-Associated Fibroblasts; PVL, Perivascular-like cells. **a** Output of cell-type deconvolution using DestVI after preprocessing. **b** Spatial map of predicted single-cell types obtained with HEDeST, revealing fine-grained tissue organization. **c** Representative mosaics of predicted cell types, showing characteristic morphologies consistent with known cellular phenotypes. **d** Distribution of nuclear areas across predicted cell types (amongst a total of 45,237 cells). **e** Spot-level co-localization matrix showing Pearson correlations between spot-wise cell-type proportions. **f** Mean inter-cell-type distance matrix derived from neighboring cells identified through Delaunay triangulation. **g** Cell-level co-localization graph summarizing neighborhood composition; arrow width indicates the average frequency of one cell type among another’s neighbors, highlighting strong co-localization among immune populations. **h** Neighborhood composition clustering. Cells were grouped based on the composition of their local neighborhoods, revealing biologically interpretable clusters. The resulting clusters correspond to coherent and biologically interpretable tissue structures, consistent with observable histological organization. Scale bar indicates 50 µm.

We next explored whether HEDeST could reveal co-localization patterns. Classical colocalization analyses rely on correlations between spot-level proportions [44] (**Fig. 5e**). In contrast, HEDeST enabled us to construct a cell-level spatial graph (see Methods), allowing the study of single-cell neighborhoods. This allowed us to investigate average inter-cell-type distances (**Fig. 5f**) and neighbor counts (**Fig. 5g**), suggesting stronger interactions among immune cells. To capture specific spatial motifs in cellular arrangements, we adopted a neighborhood-composition approach: for each cell, we quantified the cell type proportions in the direct neighborhood. Then, we clustered cells based on these neighborhood composition vectors using a Gaussian Mixture Model (GMM). This analysis identified eight neighborhood composition clusters (**Supp. Fig. 23**), two of which displayed highly similar profiles and were therefore grouped together, yielding seven well-defined clusters (**Fig. 5h**).

Cluster 0 was composed exclusively of epithelial tumor cells, characterized by a disorganized proliferation of high-grade, nucleolated cells consistent with DCIS displaying a solid architectural pattern. The identification of this cluster is clinically significant, as such high-grade lesions are frequently associated with micro-invasive foci, requiring careful histological examination of the surrounding stroma to rule out invasive carcinoma. Indeed, in several regions of the Tissue Micro-Array (TMA), Cluster 0 exhibited ambiguous architectural patterns where the boundary between the solid DCIS and the adjacent stroma appeared blurred; these ‘equivocal’ zones represent high-priority areas that would typically require complementary immunohistochemistry such as p63 to assess the integrity of the myoepithelial cell layer and formally exclude micro-invasive progression.

At its periphery, Cluster 5 circumscribed these carcinomatous masses, acting as a distinct spatial interface between the tumor and the surrounding stroma. This cluster was predominantly composed of epithelial tumor cells but also integrated a significant microenvironmental component, including endothelial cells, cancer-associated fibroblasts (CAFs), and myeloid cells. The recruitment of these diverse cell types at the boundary suggests that Cluster 5 captures the active tumor-stroma interface, where the interplay between malignant cells and the remodeled stroma facilitates metabolic exchange and immune modulation [45, 46].

Mainly localized on the left edge of the TMA, Cluster 1 was primarily composed of T cells, alongside substantial proportions of B cells, myeloid cells, and scattered plasmocytes. Its close spatial proximity to the tumor margins is highly consistent with the formation of tertiary lymphoid structures (TLS), which serve as local hubs for anti-tumor immune coordination [47, 48]. According to the maturation framework proposed by Guillaume et al. [49], the absence of a clearly defined follicular organization or active germinal centers suggests that these structures remain in an immature or ‘early-stage’ state. Unlike mature TLS, these aggregates likely represent primary lymphoid recruitment sites that have not yet achieved full functional compartmentalization.

Clusters 2 and 3, which were found ubiquitously distributed throughout the stroma, were predominantly composed of plasmocytes, CAFs, and myeloid populations such as macrophages and monocytes, reflecting a diffused inflammatory and fibroblastic landscape surrounding the tumor domains.

Together, these findings show that HEDeST enables a higher-resolution dissection of the TME. By leveraging both morphological features and spot-level priors, it reveals biologically coherent domains of cellular organization that remain inaccessible to classical deconvolution approaches (**Supp. Fig. 24**). However, despite the clinical significance of central comedonecrosis, a marker of poor prognosis characterized by ghost cells and karyorrhectic debris, our method did not distinguish this necrotic compartment from viable tumor tissue. Similarly, hydroxyapatite microcalcifications were not resolved as distinct spatial entities. These limitations are primarily due to their acellular nature: because these regions lack intact nuclei, they were not segmented by HoVerNet during the initial processing stage. Furthermore, their mineral or debris-based composition lacks the transcriptomic signal required for mapping, preventing them from being identified as separate biological domains within the HEDeST framework.

## 3 Discussion

In this study, we introduced HEDeST, a weakly supervised framework for single-cell type annotation directly from histology images, guided by deconvolution-derived spot-level proportions. Our results demonstrate that HEDeST can reliably recover cell-type identities at single-cell resolution, including for cells located outside capture spots, and that it outperforms existing alternatives in both simulated and semi-simulated settings. By combining morphological embeddings with spot-level priors and refining predictions through PPSA, HEDeST bridges the gap between ST-based deconvolution and morphology-based annotation, providing an interpretable and scalable tool for spatial biology.

A key strength of HEDeST is its flexibility. Unlike pretrained histology models, which are constrained to fixed categories and often suffer from batch effects when applied across different datasets [50, 51], HEDeST performs slide-specific training. This ensures robustness to technical variability and enables the user to define the exact set of cell types of interest, matching those in the deconvolution reference. Furthermore, because the framework can be paired with any deconvolution method, it integrates seamlessly into existing analysis pipelines and will naturally benefit from ongoing methodological advances in deconvolution algorithms. Our benchmarks also demonstrate the strengths of HEDeST in multiple contexts. Compared to HistoCell [27], HEDeST achieved higher single-cell classification accuracy, which can be attributed to its architecture and the incorporation of PPSA. In comparison with HoVerNet [28], HEDeST benefits from slide-specific training and can predict a broader, customizable set of cell types.

HEDeST offers multiple avenues for biological discovery and practical utility. By providing celllevel annotations, it enables a finer characterization of tissue organization, allowing the study of spatial neighborhoods and cell co-localization that are otherwise inaccessible to deconvolution alone. A key future perspective involves leveraging HEDeST’s single-cell resolution to achieve even finer characterization of cell populations within complex tissue interfaces, such as the tumor–stroma boundary. By refining cellular identities in these regions, the framework could help resolve “equivocal” zones where boundaries between DCIS and invasive carcinoma blurs. This enhanced granularity would enable more precise mapping of cellular and extracellular matrix remodeling, offering deeper insights into early tumor invasion and disease progression. HEDeST also represents a scalable alternative for generating large-scale single-cell annotations with minimal manual input. Moreover, this framework enables ST-informed reanalysis of histology data by providing high-resolution cell-type annotations that can be leveraged for downstream studies of clinical outcomes, treatment response, or other clinically relevant endpoints.

Despite these strengths, our study also has limitations. First, HEDeST does not integrate transcriptomic and morphological modalities in a fully joint manner. Instead, it relies on deconvolution-derived priors as the guiding signal and morphology as the predictive feature space. In principle, integrating both modalities within a unified model could further enhance accuracy and robustness [14]. On the other hand, this design has also advantages in that it can be applied on top of existing deconvolution algorithms and therefore be applied to existing studies in a post-analysis. Second, HEDeST inherits limitations from the deconvolution step, so inaccuracies in the primary deconvolution can propagate to HEDeST’s predictions. Nevertheless, the modular approach that we propose allows HEDeST to benefit from novel developments in deconvolution. Third, platform-specific artifacts such as RNA diffusion in Visium [52] or variable tissue quality can constrain both the priors and the morphological features, ultimately limiting performance of any algorithmic approach. Finally, while HEDeST adapts well to single slides, scaling to very large cohorts may require efficient single-nucleus encoders, robust to slide-level variations in staining and scanners, thus improving computational efficiency and robustness.

Looking forward, these limitations suggest natural extensions. Developing joint models that directly integrate image and transcriptomic signals could reduce dependency on deconvolution accuracy and capture complementary information from both modalities. The downside of such an approach is a certain lack of flexibility, as provided by HEDeST, but it remains to be seen whether the performance gains could potentially outweigh this drawback. While HEDeST already provides cell-level probability estimates, formalizing uncertainty modeling, such as distinguishing model-driven versus data-driven uncertainty or calibrating confidence intervals, could further enhance interpretability in clinical contexts. Moreover, HEDeST should also be applicable to other ST technologies and could be extended to higher-resolution platforms like Visium HD, where deconvolution is still necessary to resolve mixed-cell bins [13].

## 4 Conclusions

HEDeST provides a practical and effective solution to bridge the resolution gap between ST and histology-based analysis, achieving accurate single-cell annotations that are robust to technical variation and adaptable to user-defined cell types.

Overall, HEDeST represents a step toward more comprehensive tissue profiling, offering a flexible framework for integrating histology and transcriptomics. By refining cell-type resolution and revealing spatial interactions, it has the potential to advance our understanding of tissue organization in health and disease. As sequencing and imaging technologies continue to advance, frameworks like HEDeST will be key in making spatial biology useful for cancer research and clinical applications.

## 5 Methods

### 5.1 Data collection and preprocessing

Xenium data. Xenium data were downloaded from the 10x Genomics website [34, 35]. Xenium is an imaging-based spatial transcriptomics platform that provides subcellular-resolution gene expression measurements directly in tissue sections and typically includes cell segmentation and spatial coordinates. The raw data were provided in compressed format and converted into a Zarr object to ensure compatibility with the spatialdata Python library [53], which was used for downstream preprocessing and analysis. The original datasets contained 167,780 cells and 313 genes for the Human Breast Cancer slide, and 162,254 cells and 377 genes for the Human Lung Cancer slide. No specific preprocessing was performed on the Human Breast Cancer slide. For the Xenium Human Lung Cancer slide, only genes shared with the Lung Cancer Atlas [54, 36] were retained, and cells with total expression counts below 10 were removed to exclude low-coverage cells, resulting in 139,512 cells and 174 genes after preprocessing.

Visium data. The Visium Human Breast Cancer (FFPE) slide was retrieved from the 10x Genomics website [39]. The original dataset contains expression of 17,943 genes across 2,518 spots (**Supp. Fig. 21**). Preprocessing with Scanpy library [55] included removing spots with more than 20% mitochondrial content; mitochondrial genes were identified via the ‘MT-’ prefix. Genes were filtered using the *‘filter_genes’* function with the parameter *min_cells=10*. After intersection with the preprocessed scRNA-seq data (see below), the number of genes in the ST data was reduced to 2,668. Across spots, a minimum of 53, an average of 771, and a maximum of 1,298 genes had non-zero expression; total expression per spot ranged from 322.59 to 3,119.69 (mean 2,209.43). After segmentation, 45,237 cells were counted.

#### Human Lung Cancer Atlas

The Lung Cancer scRNA-seq dataset [54] was retrieved from cellxgene website [36]. This dataset contains more than 1.2 million single cells derived from 19 studies and 309 patients. We used the general annotations provided in the dataset, corresponding to the ‘ann coarse’ label, to define broad cell type categories across samples. The data were preprocessed with Scanpy: gene expression counts were normalized to constant library size across cells and log-transformed. Highly variable genes were then identified using thresholds of *min_mean=0.0125, max_mean=3*, and *min_disp=0.5*. The dataset was restricted to these highly variable genes for downstream analyses. It was also restricted to genes shared with the Xenium Lung Cancer dataset to ensure consistency. Cells with zero expression across these shared genes were removed from the analysis.

Human Breast Cancer Atlas. The Breast Cancer scRNA-seq dataset [40] was obtained from the Single Cell Portal (Broad Institute) [41]. The dataset contains a total of 100,064 cells, which were annotated according to the ‘celltype_major’ label, corresponding to nine major cell types. Cells annotated as Normal Epithelial were excluded from the analysis. According to our pathologist, the cell type originally labeled as Plasmablasts in the atlas was renamed to Plasmocytes, as this designation better reflects the cell types typically observed in breast cancer tissues. Quality control and preprocessing were performed with the Scanpy library. We filtered out cells with more than 100,000 total counts, more than 8,000 detected genes, or a mitochondrial fraction greater than or equal to 20%. At the gene level, we filtered out genes with fewer than 10 total counts. Raw counts were preserved in a counts layer, HVGs were identified with the Seurat v3 method, and the top 4,000 HVGs were retained. After preprocessing and intersection with ST data, the single-cell reference consists of 95,624 cells and 2,668 genes.

Histology data. The H&E images were retrieved together with the ST data from the 10x Genomics website. The files are provided in TIFF format for Visium data and in OME-TIFF format for Xenium data. The histological images were further preprocessed to generate tissue masks, which is a crucial step before performing segmentation. For this purpose, we implemented an automatic masking function that creates a binary mask from a downsampled version of the WSI. The process involves converting the RGB image to grayscale, removing potential border artifacts, and applying Otsu’s thresholding method [56] to distinguish tissue regions from the background. To refine the mask, morphological operations such as closing and opening are applied, which help remove noise and small irregularities. The resulting binary mask highlights the tissue area while suppressing irrelevant background, thereby ensuring that subsequent segmentation analyses focus only on biologically relevant regions of the histological slide.

### 5.2 HEDeST method

#### H&E image segmentation and tile extraction

Nucleus segmentation is performed with HoVerNet [28], using the pre-trained weights *‘hovernet fast pannuke type tf2pytorch*.*tar’* available in their Github repository. The entire WSIs are processed using the *run wsi*.*sh* script and default parameters. This step provides segmentation information for each nucleus, including contours and centroids, along with classification into five broad categories (neoplastic, non-neoplastic, necrosis, connective, and inflammatory). From these results, we cropped 20 µm × 20 µm single-cell image patches centered on each nucleus centroid, which were subsequently resized to 64 × 64 pixels. We selected a 20 µm crop because this size yielded the highest predictive performance and fully captures larger cells. For smaller cells, the crop can include additional surrounding context beyond the nucleus, which may be suboptimal, yet no single crop size provides an ideal compromise across all cell types.

#### Self-supervised learning

In order to obtain a robust representation of cells, we use a contrastive learning approach [57]. This family of methods aims to bring together, in the representation space, different images of the same cell (or similar cells) while distancing those of distinct cells, thus allowing the model to learn discriminating features without supervision. For this purpose, we have selected the moco-v3 model from facebook [29]. The architecture chosen is a ResNet-50, trained for 150 epochs with a batch size of 2048. At the end of training, each cell is represented by a 2048-dimensional embedding vector, which captures the morphological information relevant for further analysis.

#### Learning from label proportions

We define the following variables:

- *N*_*c*_: total number of cells,
- *N*_*s*_: total number of spots,
- *N*_*t*_: number of cell types,
- *n*_*i*_: number of cells contained in spot *i, i* ∈ {1, …, *N*_*s*_},
- 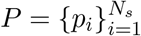: matrix of proportions with dimensions *N*_*s*_ × *N*_*t*_, obtained from the deconvolution of ST data,
- 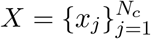: matrix of cell embeddings with dimensions *N*_*c*_ × 2048, where each vector *x*_*j*_ represents the morphological features of cell *j*,
- *p*(*t*|*x*_*j*_): probability vector representing the likelihood of cell *j* belonging to each of the *N*_*t*_ cell types.

HEDeST is based on LLP [30], a weakly supervised learning approach where individual labels are not available, but only class proportions are known at the group level. In our case, the cell type proportions per spot serve as supervision to train a cell-level classifier.

We train a classifier *f* (*x*_*j*_) = *p*(*t*|*x*_*j*_), implemented as a multilayer perceptron (MLP) that takes as input a 2048-dimensional cell embedding and outputs a probability distribution over *N*_*t*_ cell types. The architecture consists of two successive linear layers of sizes 512 and 256, each followed by a layer normalization and a ReLU activation, and a final linear layer projecting into ℝ^*N*^t, followed by a *softmax* function (**Supp. Fig. 1**). Since cell-level ground truth annotations are not available, training relies on aggregation at the spot level. For each spot *i*, the predictions of its constituent cells are averaged:

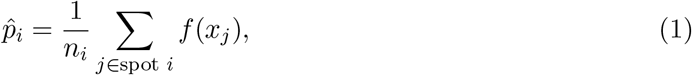

where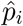 denotes the predicted cell-type proportions in spot *i*. Cells outside of any spot are excluded during training.

Because training is supervised by spot-level proportions, the loss function is computed independently for each spot and then averaged across the spots within a batch during optimization. The loss for a given spot *i* is defined as the mean squared error (MSE) between the predicted proportions 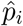and the reference proportions *p*_*i*_:

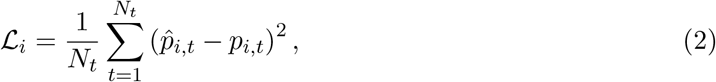

where *p*_*i,t*_ and 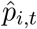denote the observed and predicted proportions of cell type *t* in spot *i*, respectively.

The model is trained for 100 epochs with a batch size of 64 and a learning rate of 1 × 10^−4^, using a dataset split into 80% for training, 10% for validation, and 10% for testing. Optimization is carried out using the Adam optimizer, and the model is implemented in PyTorch (version 1.13.1+cu116) with explicit CUDA support, allowing computations to be executed on GPUs for efficient large-scale training.

After training, the model weights corresponding to the lowest validation loss are frozen. The model is then applied to all cells in the H&E slide, including those outside the spots, to infer their probability vectors *p*(*t*|*x*_*j*_). The final output of HEDeST is therefore a matrix of dimension *N*_*c*_ × *N*_*t*_, where each row corresponds to the probabilistic profile of a single cell across the considered cell types. In downstream analyses, the attributed cell type of each cell is defined as the class associated with the maximum probability.

#### Prior Probability Shift Adjustment

In the output of the deconvolution step, the cell type proportions satisfy

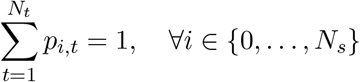

which allows us to interpret *p*_*i*_ as a localized prior probability distribution of cell type occurrence in spot *i*. From model training, we obtain for each cell *j* the posterior probability distribution *p*(*t*|*x*_*j*_). According to Bayes’ theorem, the posterior probability can be written as:

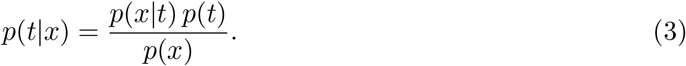

Here, all probabilities are estimated from the training set. However, it is important to note that in a given spot, the local prior distribution may differ substantially from the global prior in the training set. To account for this discrepancy, we replace the global prior with the spot-specific proportions obtained from deconvolution. We therefore define a corrected posterior probability:

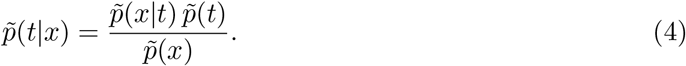

We make the assumption that the class-conditional data densities remain unchanged between the global training set and the local spot, i.e.:

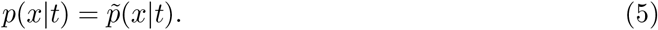

Under this assumption, the corrected posterior probability can be expressed as:

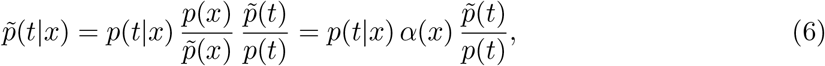

where 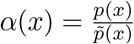is a normalization factor. It can be determined by enforcing that the corrected posterior probabilities sum to one:

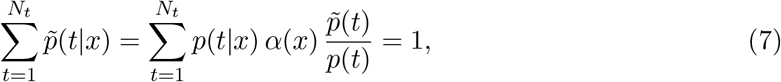

which yields the expression:

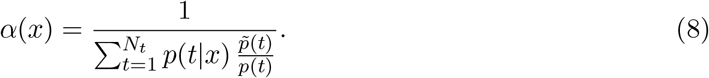

Finally, substituting *α*(*x*) into Equation (7), we obtain the corrected posterior distribution in its practical form:

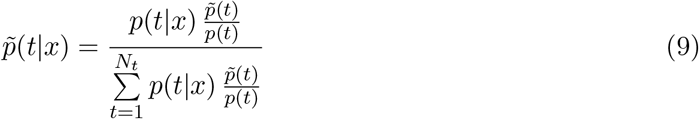

This expression corresponds to a reweighted version of the original posterior. Of note, the training set prior probability *p*(*t*) can be interpreted as the global average proportion vector across the training set, i.e.

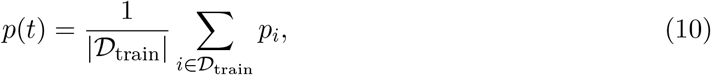

whereas the spot-specific prior corresponds directly to the local proportion :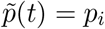.

#### Spatial Adjustment of cells outside spots

Cells that fall outside spots lack a direct estimate of their local cell-type composition. To enable prior probability shift adjustment in this setting, we construct for each such cell a synthetic local prior distribution 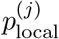by averaging nearby spot-level proportion vectors. For cells located inside a spot, the local prior is directly given by the spot composition itself. Formally, if cell *j* belongs to spot *i*, then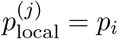.

For every cell outside spots, we query its centroid position against the set of spot centroids. The neighbourhood consists of the nearest *k* ≤ 3 spots located within a search radius *R* = 2 × spot diameter. If no spot lies within this radius, the cell is deemed unadjustable and retains its uncorrected probability vector.

When at least one neighbouring spot is found, its contribution to the local prior is weighted according to spatial proximity. For a neighbour at distance *d* from the cell, the weight is defined as

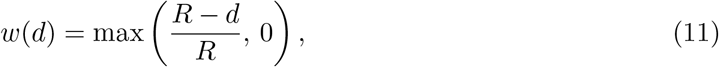

If multiple neighbours are available, their proportion vectors are combined by a normalized weighted average:

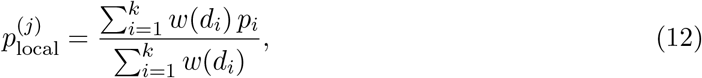

In the special case where exactly one neighbour is within range, a trade-off coefficient *β*_*j*_ is made distance-dependent. The intuition is that a cell directly overlapping a spot should adopt the neighbour’s prior almost entirely (*β*_*j*_ ≈ 0), while a cell further away should rely more strongly on its original prediction (*β*_*j*_ ≈ 1). The coefficient is computed as

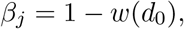

where *d*_0_ is the distance to the single neighbour. For other cells, *β*_*j*_ is set to 0 by default. Now that we have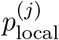 for all cells, we perform PPSA using it as the local prior. Concretely, 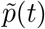can be replaced by 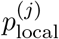 in Equation (9). Finally, for each cell, the adjusted prediction is interpolated with the original output using the coefficient *β*_*j*_:

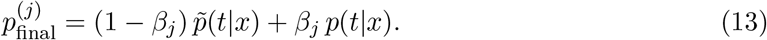

#### Downstream analysis

We have developed several tools to visualize and exploit HEDeST annotations. These tools include interactive or static visualizations of annotated cells on H&E images, as well as galleries showing mosaics of cells for each cell type and spot-by-spot visualizations. Our analysis functions allow us to extract basic descriptive statistics, evaluate the quality of the results, and compare them with a ground truth, if available.

A significant part of our analyses is based on the study of cell neighborhoods. To this end, we have implemented a Delaunay graph, which consists of a triangular mesh connecting the nearest cells to each other [58]. This graph allows us to study cell-cell or cell type-cell type interactions, providing a structured framework for analyzing spatial relationships within the tissue.

### 5.3 Simulations

#### Fully simulated data

To generate a fully simulated dataset, we used the H&E image from the Visium FFPE Human Ovarian Cancer dataset [31]. We first performed nucleus segmentation and self-supervised learning to extract cell embeddings, then applied clustering using the K-means method (Euclidean distance). In order to obtain morphologically distinct clusters, we retained only the 2,000 cells closest to the centroid for each cluster. The remaining cells were then assigned to bags, hereafter referred to as ‘pseudo-spots’, according to a normal distribution parameterized by a mean (average number of cells per spot) and a variance. Depending on the experiments, we varied those parameters but also the overall fraction of each cluster in the set of pseudo-spots. For balanced simulations, all clusters were equally represented across pseudo-spots. For imbalanced scenarios, overall cluster proportions were set to [0.5625, 0.0625, 0.125, 0.25] for clusters 0–3 in the 30-spot simulation, and [0.1, 0.0143, 0.6286, 0.0571, 0.0143, 0.1286, 0.0571] for clusters 0–6 in the 200-spot simulation. Finally, we calculated the pseudo-spot-specific proportion vectors describing the composition in clusters.

#### Semi-simulated data

We worked with two Xenium datasets for which nuclei and cells had been pre-segmented based on RNA distribution. For the Xenium Human Breast Cancer dataset [33, 34], nuclei were provided with annotation covering 19 distinct cell types, which we grouped into 10 broader classes. For the Xenium Human Lung Cancer dataset [35], we performed cell type annotation using a scRNA-seq reference [36]. Specifically, for each Xenium cell, we identified its 20 closest neighbors in the reference dataset. As a metric, we chose cosine distance as it emphasizes similarity in expression profiles by comparing vector orientation rather than magnitude, making it robust to differences in sequencing depth. Each Xenium cell was then annotated based on the majority cell type among its 20 closest neighbors [59]. Finally, Xenium cells were matched to Xenium nuclei, keeping only associations with more than 95% overlap between nuclear and cellular masks.

We aligned the H&E image, previously segmented with HoVerNet, to the Xenium data using the spatialdata library. We then matched the Xenium nuclei with the HoVerNet-segmented nuclei. Since slight coordinate shifts between modalities sometimes prevented direct overlap, matching was performed based on the closest nucleus within a maximum distance of 40 pixels (**Supp. Figs. 6b, 12b**).

Following this procedure, most of the nuclei segmented by HoVerNet inherited the cell type annotation of their corresponding Xenium nuclei and were used for semi-simulation. Nuclei could not be annotated in cases where (i) the number of RNA counts was too low to allow for reliable annotation, (ii) no corresponding nucleus or cell was identified during the matching steps, or (iii) the assigned cell type was rare, defined as fewer than 1,000 cells across the whole slide, in which case both the nuclei and the corresponding cell type were removed. We then created our pseudo-spots by generating grids of circular regions, respecting real Visium spot layout, which were overlaid on the H&E slide. Each cell whose nuclear centroid was located within a pseudo-spot was assigned to it, while cells located outside remained unassigned. Since the cell type of each cell was known, we computed cell type proportion vectors within each pseudo-spot.

#### Experiments with proportion perturbation

To evaluate the robustness of HEDeST to inaccuracies in spot-level priors, we introduced controlled perturbations to the simulated proportion matrices. Specifically, for each pseudo-spot, we combined the original cell-type proportions with randomly generated noise, using a mixing parameter that defines the perturbation level. A strength of 0 corresponds to the unaltered proportions, while a value of 1 replaces them entirely with random values. After adding noise, proportions were re-normalized to ensure that each spot’s vector still summed to 1. This approach allowed us to systematically vary the reliability of the priors and assess HEDeST’s performance under different degrees of proportion disruption.

### 5.4 Benchmark study

#### MHAST

MHAST [32] combines cell-type deconvolution with morphological features to assign cell types to cells. First, it estimates cell type counts per spot using deconvolution methods such as Tangram. In parallel, nuclei are segmented directly from the H&E image and morphological features are extracted. To resolve the correspondence between deconvolved cell type counts and individual nuclei, MHAST formulates the task as an optimization problem on possible label assignments and achieves it through hierarchical permutation.

MHAST was applied to simulated data consisting of 4 morphological clusters distributed across 30 pseudo-spots. These values were selected since the optimization problem scales exponentially with the number of clusters and spots, making larger configurations computationally infeasible. The number of cells per pseudo-spot followed a normal distribution with a mean and variance both equal to 5. We evaluated two scenarios: one in which all clusters were equally represented across the pseudo-spots, and a more realistic one characterized by a strong imbalance between clusters (**Figs. 2a, 2b**).

#### HoVerNet

We retrieved the broad cell type predictions made by HoVerNet on the Xenium slides. Cells labeled as necrosis or non-neoplastic were excluded due to their low abundance. To ensure comparability, we reannotated the semi-simulated datasets so that HEDeST was trained and evaluated on the same broad classes (neoplastic, connective and inflammatory). Finally, we retained HoVerNet predictions for cells that had an associated ground-truth annotation.

#### HistoCell

We benchmarked our method against HistoCell [27], a deep-learning approach designed to infer cell types and cell states from histology images at single-cell level. HistoCell was originally developed as a collection of pretrained models, each trained on one of nine cancer types. Training relied on weak supervision by cell type proportions, which were obtained by deconvolving ST data using four state-of-the-art methods and averaging their results. Since the pretrained models are not publicly available, we retrained HistoCell on the semi-simulated datasets under two experimental settings: using the original scRNA-seq–derived annotations and using the 3 broad HoVerNet classes. Unlike HEDeST, HistoCell operates on 256×256 image tiles rather than individual cell images, allowing it to capture morphological features at both the cell and spot levels. Tiles were therefore extracted centered on pseudo-spots, and for each slide and experimental setting, HistoCell was trained using the pseudo-spot proportions as weak labels. Each experiment was repeated ten times with different random seeds. For inference, the entire histology slide was partitioned into non-overlapping 256×256 tiles. Each trained model was then applied to these tiles to obtain predicted cell type probabilities across the whole slide.

Because HistoCell segmentation was performed at the tile level, while ground-truth annotations were defined at the whole-slide level, the predictions were not directly comparable. To align them, we converted HistoCell cell coordinates from tile space to slide-level coordinates and matched cell contours using both centroid proximity and contour overlap, keeping only pairs with an intersection-over-union (IoU) greater than 0.3. This procedure yielded the subset of cells for which HistoCell predictions could be consistently compared with the ground truth.

### 5.5 Case study

#### Cell-type deconvolution

The Visium FFPE Human Breast Cancer slide was deconvolved using DestVI [17] with the Breast Cancer single-cell atlas [40] as reference. Briefly, the atlas was used to train the single-cell model (scLVM), which was then integrated with the ST data to fit the spatial model (stLVM). Training was performed with the ‘celltype major’ annotation as labels, a maximum of 1500 epochs for the stLVM, and default parameters otherwise. Post-processing of the deconvolution results was necessary, as the proportion of Cancer Epithelial cells was overestimated outside tumor regions (**Supp. Fig. 25**). To correct for this, Cancer Epithelial proportions between 0.2 and 0.6 were rescaled to range between 0.1 and 0.2.

Co-localization analysis. HEDeST was run on the breast cancer H&E image with cell type proportions derived from deconvolution, using default parameters. To analyze the spatial organization of cells, we first constructed a Delaunay graph in which two nodes (cells) were connected if their distance was less than 150 pixels (**Supp. Fig. 26**). We then grouped cells based on the composition of their local neighborhood. Specifically, for each cell, we calculated a vector of proportions of neighboring cell types. We then clustered the cells based on this vector using a Gaussian Mixture model (GMM). Among the eight clusters initially identified (**Supp. Fig. 23**), two were very similar, comprising a mixture of endothelial cells, CAFs, myeloid cells, and cancer cells, and were therefore merged.

To compare the cell-level clustering with a spot-level clustering, we tried two approaches. In the first approach, spots were clustered using only the cell type proportions assigned to each spot, without considering spatial context (**Supp. Fig. 24a**). In the second, spatially-aware approach, we incorporated information from neighboring spots. For each spot, we identified its six closest neighbors in physical space and constructed a row-normalized adjacency matrix representing these neighbor relationships. Each spot’s feature vector was then augmented by adding a weighted contribution of the neighbors’ cell type proportions, such that the final spatially-smoothed vector for each spot was computed as *X*_spatial_ = *X* + *λWX*, where *X* is the original spot proportion matrix, *W* is the normalized adjacency matrix, and *λ* = 0.5 controls the contribution of neighbors. The spatially-smoothed vectors were then clustered using a GMM, yielding spatially coherent spot clusters that account for both the local composition and the surrounding neighborhood (**Supp. Fig. 24b**).

### 5.6 Statistics and reproducibility

To assess model performance and compute error bars, we repeated each experiment with multiple random seeds. The resulting metrics were aggregated by calculating the mean and associated 95% confidence intervals, obtained from the standard error across runs. The variability regions on ROC curves have been obtained by interpolating the true positive rates over a fixed false positive rate grid and computing the standard deviation across seeds. In the case study, we compared the sizes of cell nuclei between different cell types using the one-sided Mann–Whitney U test. All experiments are fully reproducible, and all statistical analyses were performed using standard scientific computing libraries in Python (version 3.9.20).

## 7 Declarations

### Ethics approval and consent to participate

Not applicable.

### Consent for publication

Not applicable.

### Availability of data and materials

All codes used in this study are available in the GitHub repository at https://github.com/sysbio-curie/HEDeST. We mostly worked with data from the 10x Genomics website [39, 31, 34, 35], including Xenium Human Breast Cancer [33] (https://www.10xgenomics.com/products/xenium-in-situ/preview-dataset-human-breast), Xenium Human Lung Cancer (https://www.10xgenomics.com/datasets/preview-data-ffpe-human-lung-cancer-with-xenium-multimodal-cell-segmentation-1-standard), Visium Human Ovarian Cancer (FFPE) (https://www.10xgenomics.com/datasets/human-ovarian-cancer-11-mm-capture-area-ffpe-2-standard), from which only the H&E image was used, and Visium Human Breast Cancer (FFPE) (https://www.10xgenomics.com/datasets/human-breast-cancer-ductal-carcinoma-in-situ-invasive-carcinoma-ffpe-1-standard-1-3-0). ScRNA-seq data were downloaded from publicly available resources: the Breast Cancer Atlas [40, 41] (https://singlecell.broadinstitute.org/single_cell/study/SCP1039/a-single-cell-and-spatially-resolved-atlas-of-human-breast-cancers) and the Lung Cancer Atlas [54, 36] (https://cellxgene.cziscience.com/collections/edb893ee-4066-4128-9aec-5eb2b03f8287). All simulated data can be reproduced using the codes available in the GitHub repository.

### Author contribution

L.G, L.C., E.B, and T.W. designed and planned the study. L.G. developed the tool with support by L.C. L.G. performed the experiments, analyzed the results, prepared all figures, and wrote the manuscript. E.B. and T.W. supervised the study. R.B. provided pathological expertise, contributed to the case study, and participated in the writing of the manuscript. All authors reviewed and/or edited the manuscript before submission, and approved the final version.

## Acknowledgements

We thank H. Salmon for her expertise in cancer biology, which was useful for interpreting results. We are also grateful to L. Gaspard-Boulinc, T. Defard, and A. Díaz Herrero for their meaningful discussions and constructive feedback throughout this work.

