## Supplementary figures for "HEDeST: An Integrative Approach to Enhance Spatial Transcriptomic Deconvolution with Histology"

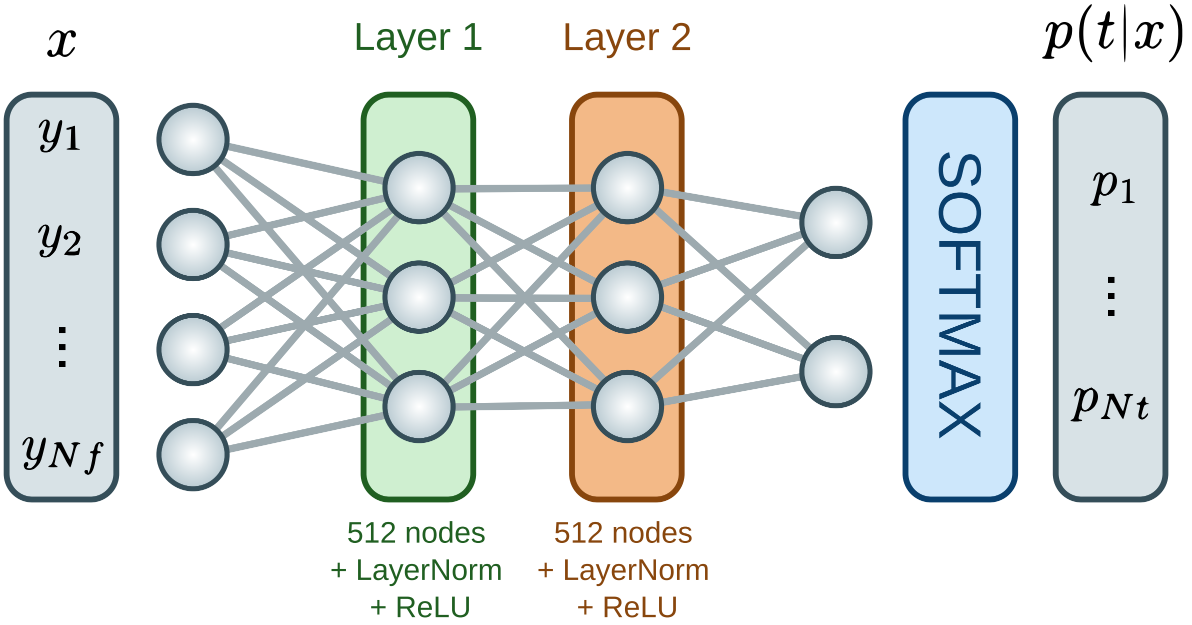

**Fig. 1** Classifier architecture used in HEDeST. The model consists of two fully connected layers with layer normalization and ReLU activation functions, followed by a final softmax layer that outputs cell-type probability estimates. Each input cell is represented by an embedding vector $x \in R^{N_f}$, where $N_f = 2048$ corresponds to the dimensionality of the MoCo-v3 features, with embedding components denoted $\{y_k\}_{k=1}^{N_f}$. The classifier produces a probability vector $p(t \mid x) = \{p_i\}_{i=1}^{N_t}$, where $N_t$ is the number of predicted cell types and $p_i$ represents the probability that the cell belongs to type $i$.

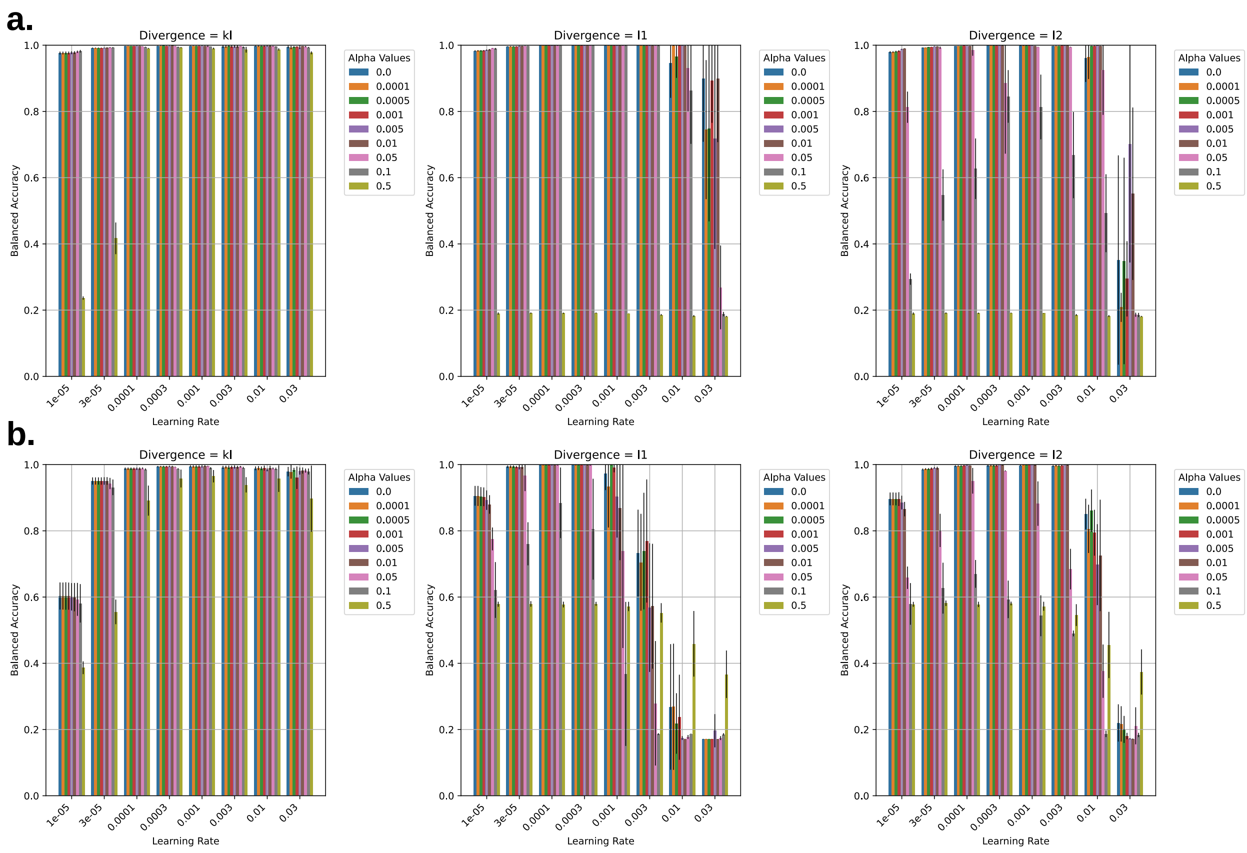

**Fig. 2** Performance plots from grid-search experiments on fully simulated data. kl, Kullback–Leibler divergence; l1, mean absolute error; l2, mean squared error. **a,b** Balanced (**a**) and imbalanced (**b**) simulation settings. We performed a grid search over loss functions, learning rates, and values of the balancing parameter $\alpha$. The original composite loss used during training for a spot $i$ was defined as:

$$\mathcal{L}_i = -\frac{\alpha}{n_i} \sum_{j=1}^{n_i} \log f(x_j)_{t_j} \; + \; (1-\alpha)\, d_K(\hat{p}_i,\; p_i)$$

where the first term encourages confident (high-probability) predictions and the second term measures the proximity between predicted and ground-truth proportion vectors through the function $d_K$ [1]. The parameter $\alpha$ controls the trade-off between these two objectives. Grid-search results showed that performance was consistently higher for $\alpha = 0$, leading us to remove the first (confidence-based) term in the final training loss.

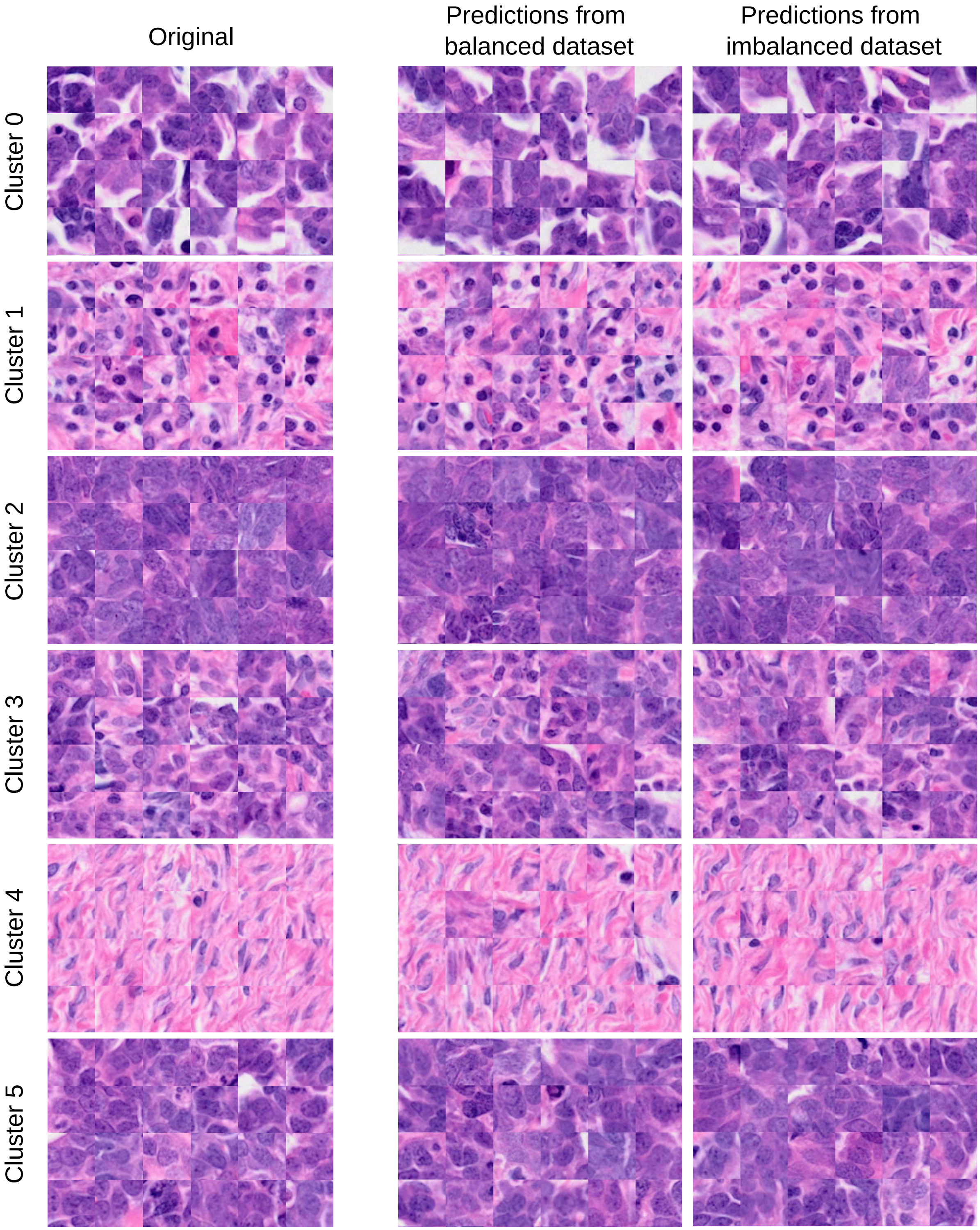

**Fig. 3** Comparison of original and predicted morphological galleries in simulated datasets. The galleries compare the morphological consistency of cell crops within each of the 6 clusters. **Left** Original galleries representing the ground-truth morphological clusters. **Center** Predicted galleries generated from the balanced simulated dataset. **Right** Predicted galleries from the imbalanced simulated dataset.

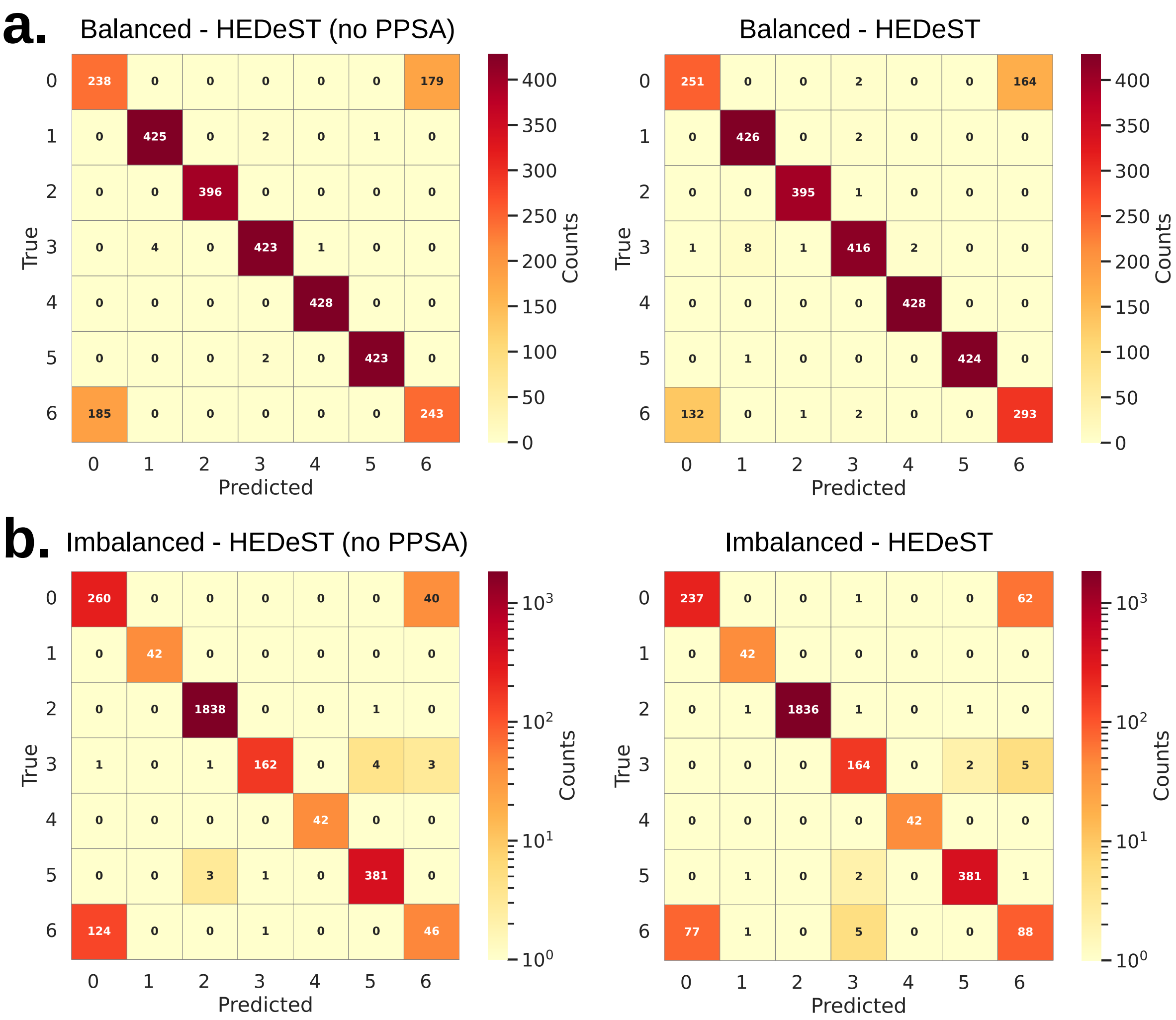

**Fig. 4** Confusion matrices for HEDeST performance on simulated datasets with duplicated clusters. HEDeST (no PPSA), HEDeST without PPSA. They evaluate classification performance on simulated datasets featuring $N_t = 7$ clusters, including a duplicated cluster. **a** Performance on the balanced dataset comparing results before (**left**) and after PPSA (**right**). **b** Performance on the imbalanced dataset under identical conditions. In both scenarios, the initial model exhibits confusion between morphologically identical clusters; the application of PPSA effectively redistributes these misclassified cells by leveraging localized proportion-based context.

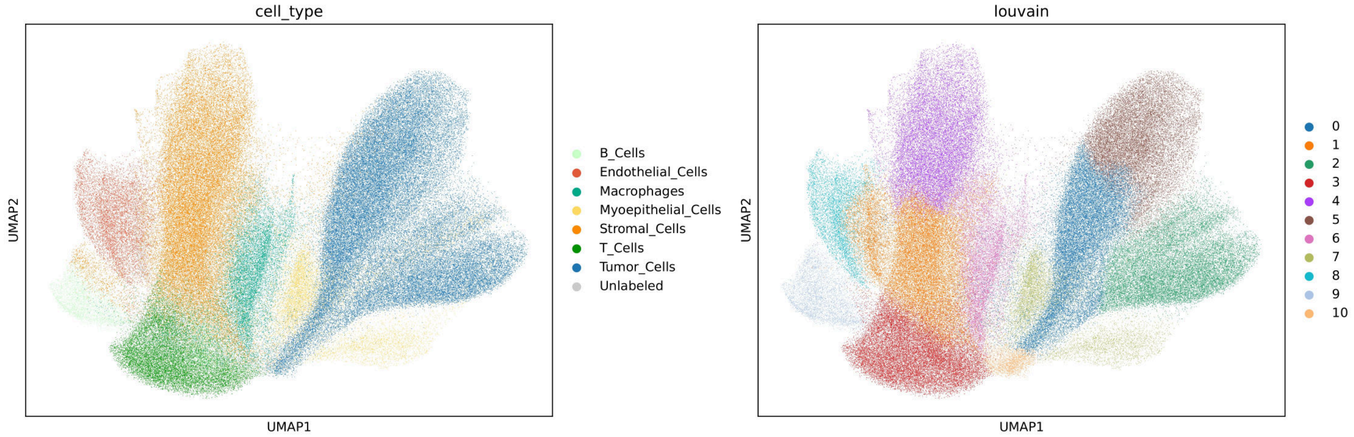

**Fig. 5** UMAP visualization of the Xenium Breast Cancer dataset. UMAP is colored by (**left**) known cell types and (**right**) Louvain clusters.

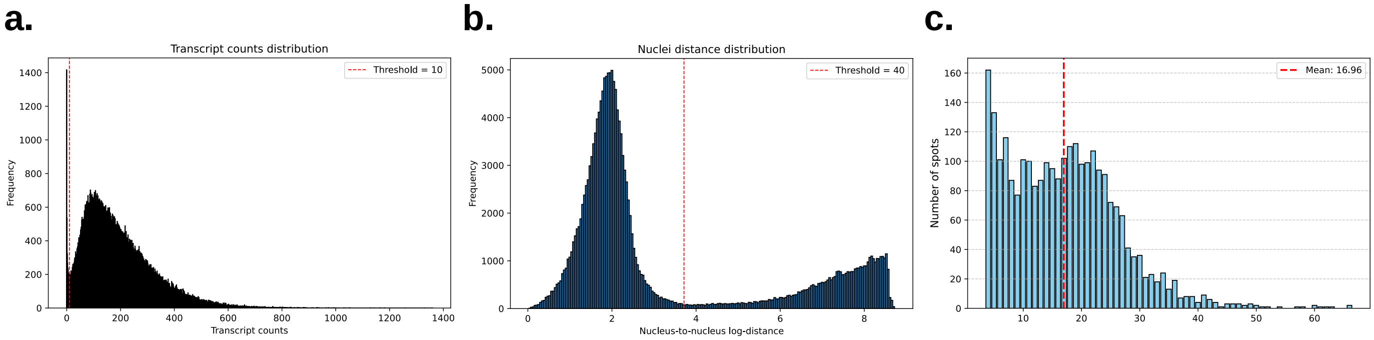

**Fig. 6** Construction details of the Xenium Breast Cancer semi-simulated dataset. **a** Distribution of transcript counts per cell (we next filtered out cells with fewer than 10 transcripts). **b** Distribution of nucleus-to-nucleus log-distances when matching Xenium and HoVerNet nuclei; only pairs with distances below 40 pixels were retained in the simulated dataset. **c** Distribution of the final number of nuclei per pseudo-spot.

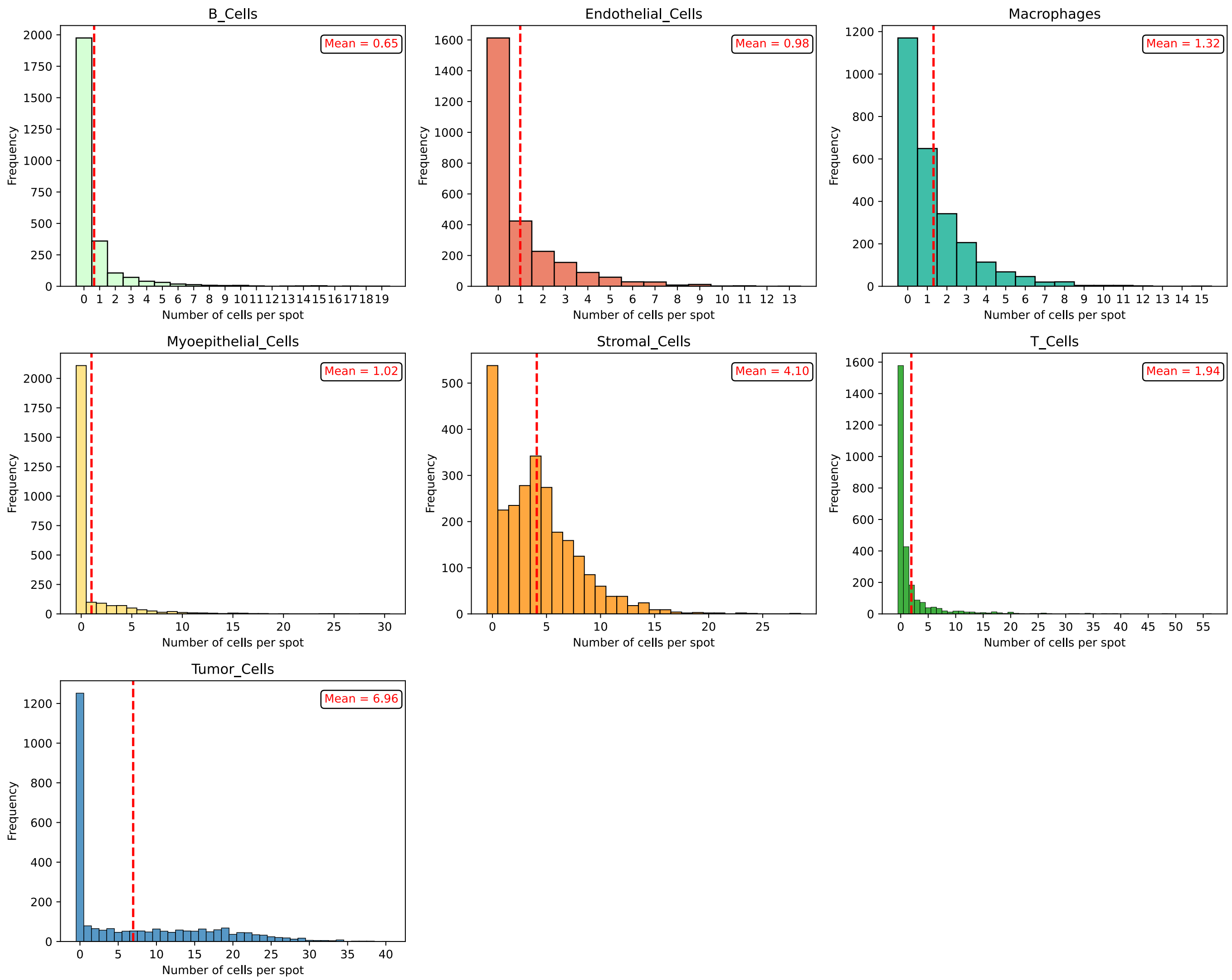

**Fig. 7** Distribution of cell types among pseudo-spots in the Xenium Breast Cancer semi-simulated dataset. The barplot shows the abundance of each cell type across all pseudo-spots, highlighting the imbalance typical of tumor tissue and the composition used for quantitative evaluation of HEDeST.

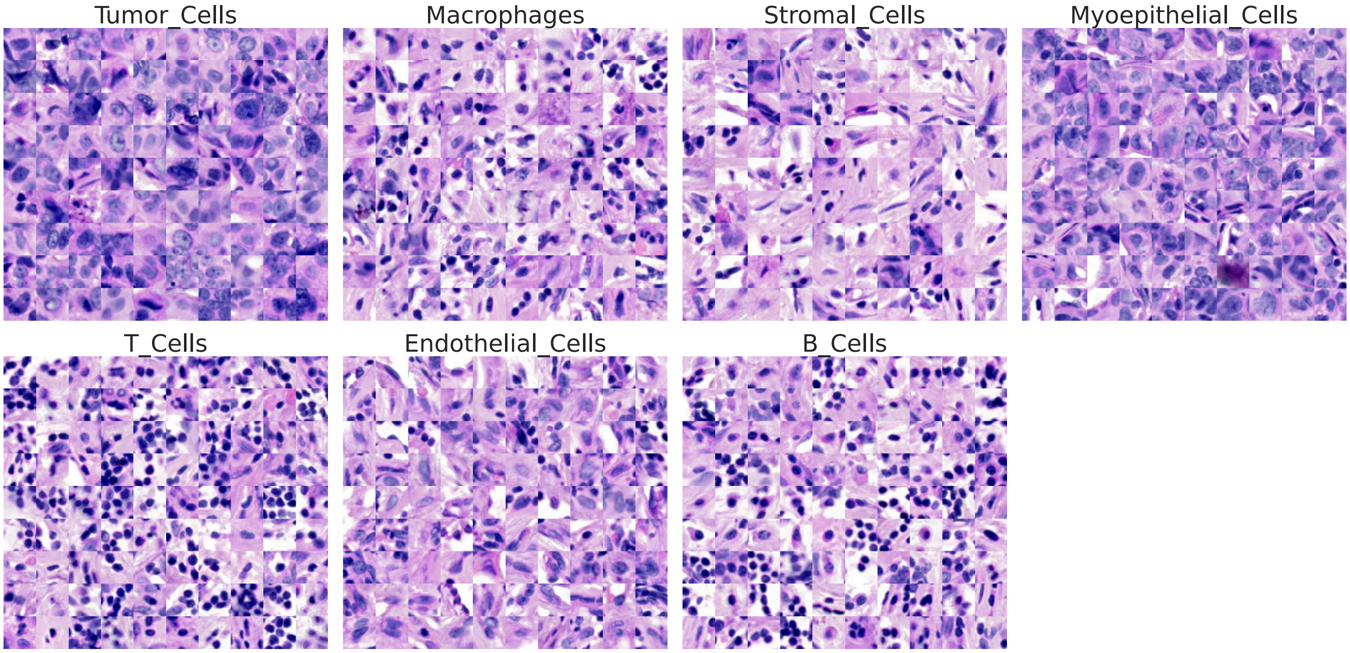

**Fig. 8** Galleries of cell type annotations in the Xenium Breast dataset.

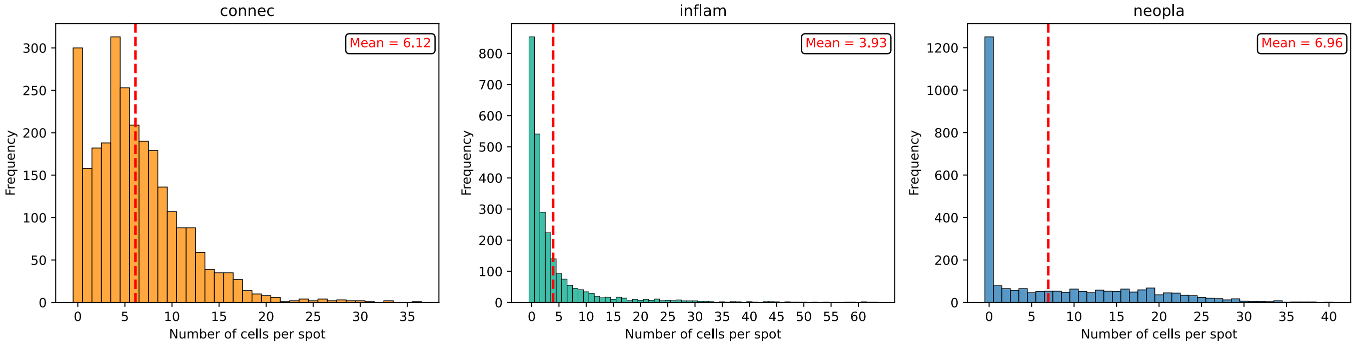

**Fig. 9** Distribution of cell types among pseudo-spots in the Xenium Breast Cancer semi-simulated dataset restricted to three broad cell types. neopla, Neoplastic cells; inflam, Inflammatory cells; connect, Connective cells.

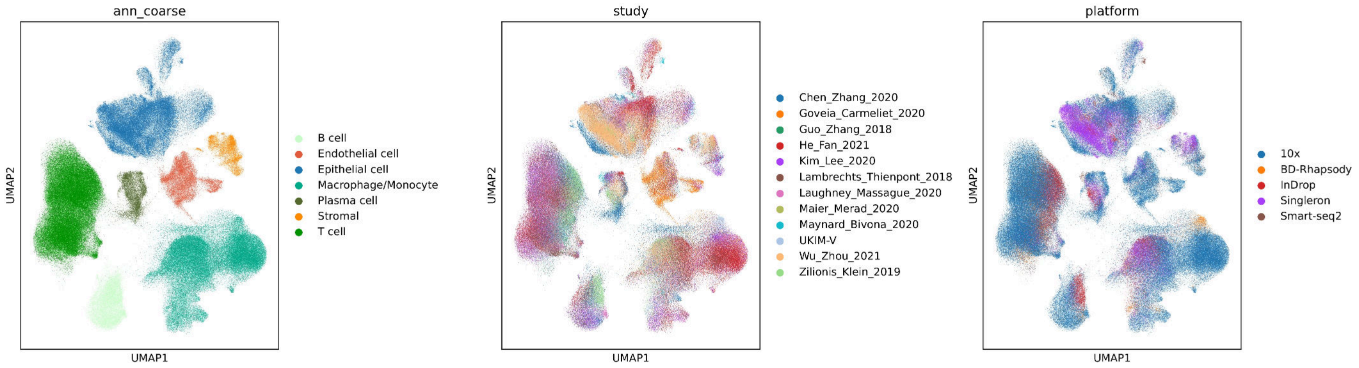

**Fig. 10** UMAP of the Lung Cancer Atlas. UMAP is colored by cell type (**left**), by study (**center**), and by platform (**right**). The projection highlights clear separation between cell types, while no clustering is observed based on study or platform, indicating minimal batch or platform effects in this single-cell reference.

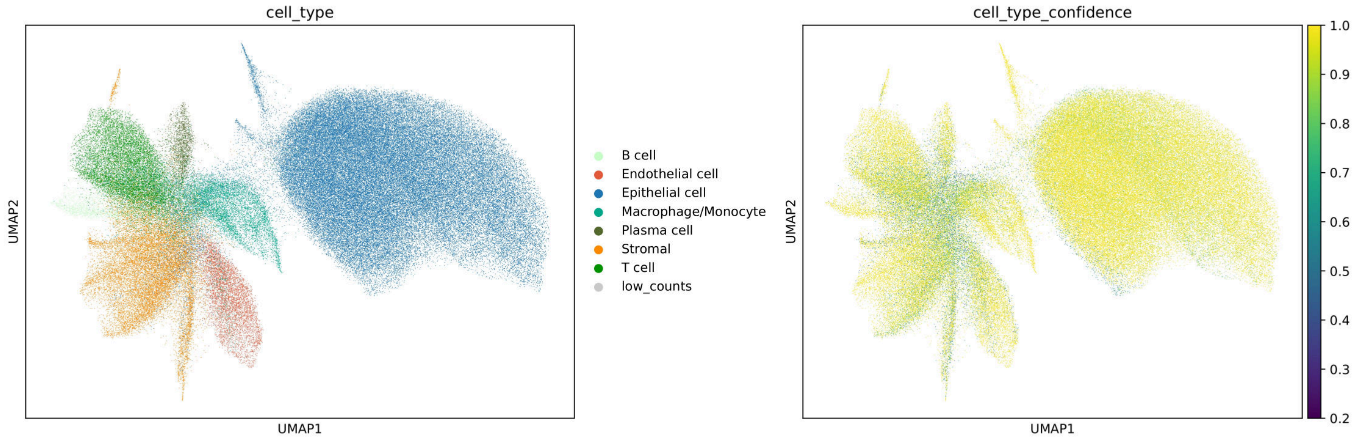

**Fig. 11** UMAP visualization of the Xenium Lung Cancer dataset. UMAP is colored by (**left**) known cell types and (**right**) annotation confidence. Annotation confidence was computed as the relative abundance of the k-nearest (k=20) single-cell neighbors (from the reference atlas) that match the final assigned cell type for each Xenium cell.

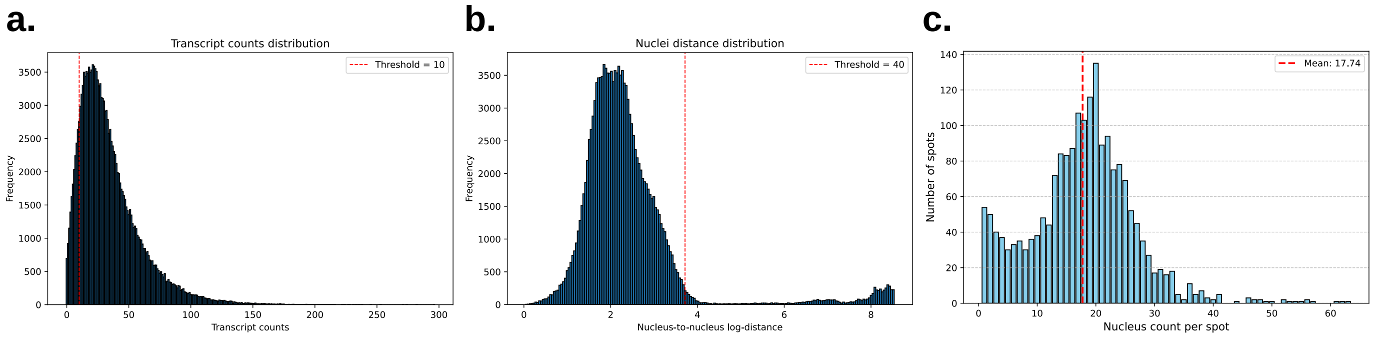

**Fig. 12** Construction details of the Xenium Lung Cancer semi-simulated dataset. **a** Distribution of transcript counts per cell. **b** Distribution of nucleus-to-nucleus log-distances when matching Xenium and HoVerNet nuclei. **c** Distribution of the final number of nuclei per pseudo-spot.

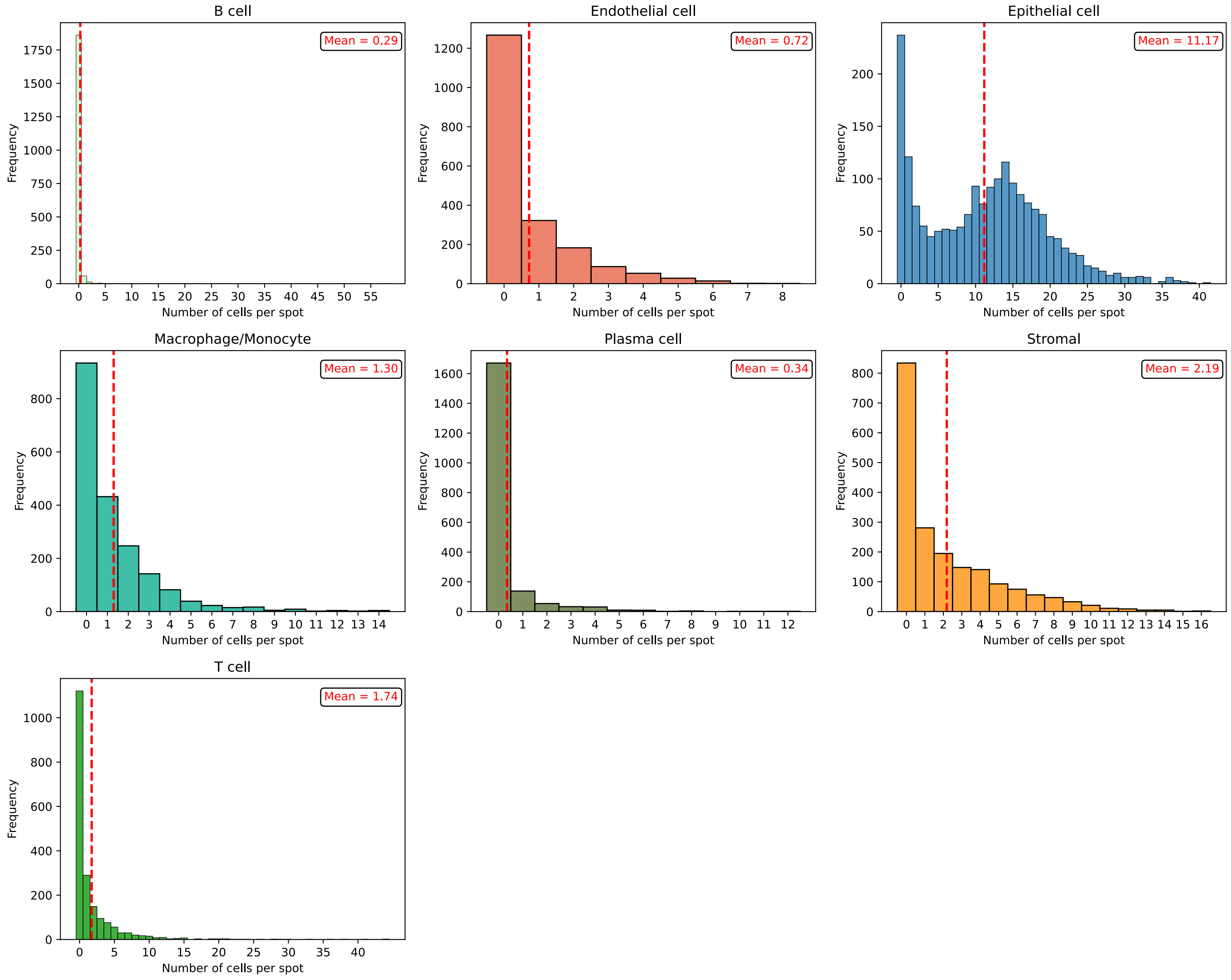

**Fig. 13** Distribution of cell types among pseudo-spots in the Xenium Lung Cancer semi-simulated dataset.

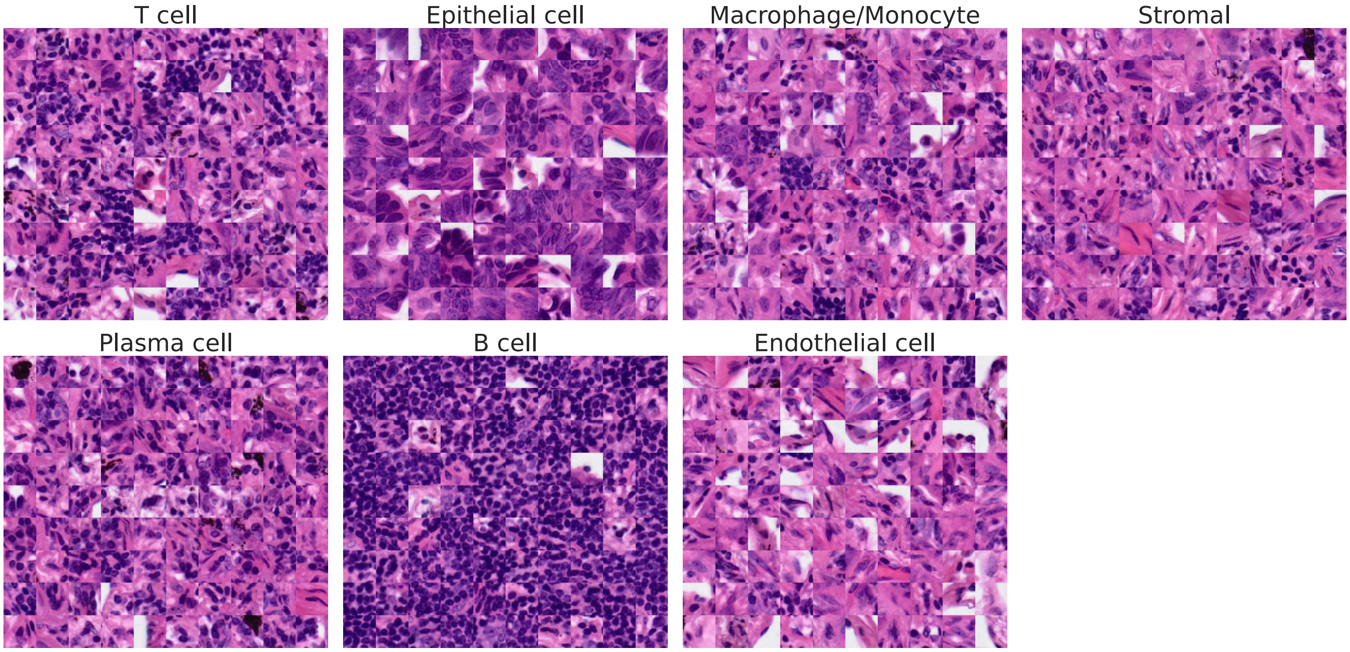

**Fig. 14** Galleries of cell type annotations in the Xenium Lung dataset.

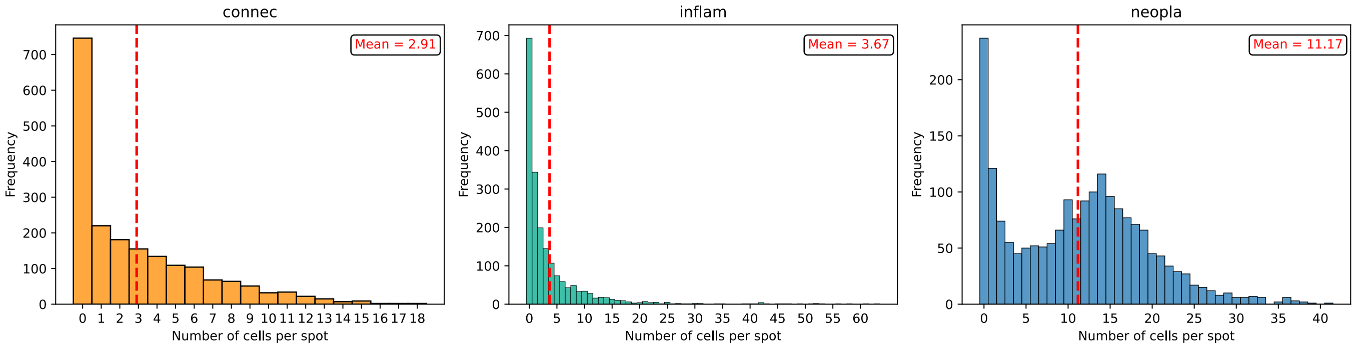

**Fig. 15** Distribution of cell types among pseudo-spots in the Xenium Lung Cancer semi-simulated dataset restricted to three broad cell types. neopla, Neoplastic cells; inflam, Inflammatory cells; connect, Connective cells.

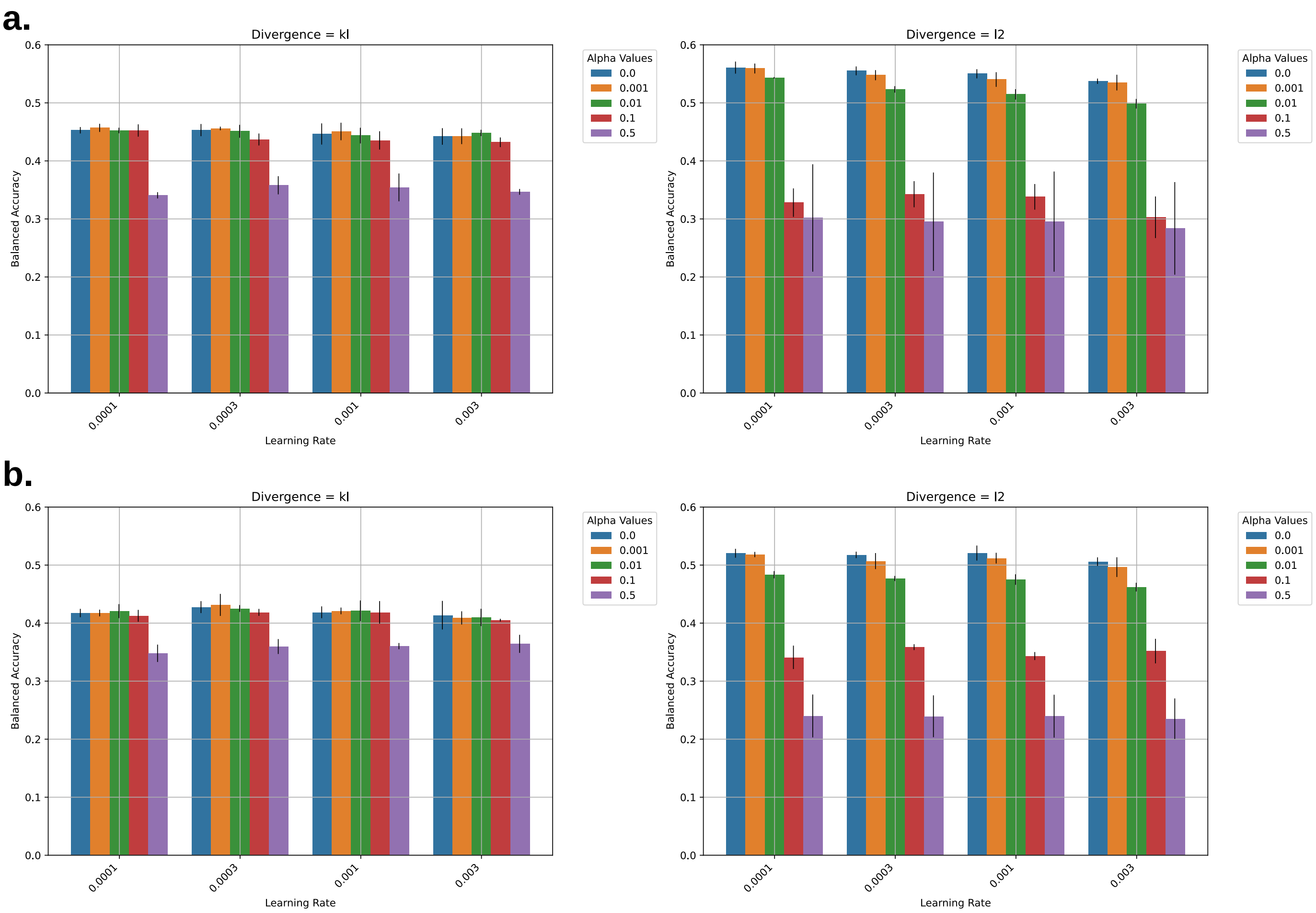

**Fig. 16** Performance plots from grid-search experiments on semi-simulated data. kl, Kullback–Leibler divergence; l2, mean squared error. **a,b** Xenium Human Breast Cancer (**a**) and Xenium Human Lung Cancer (**b**) semi-simulated datasets. Grid-search was performed over loss functions (excluding l1 loss), learning rates and values of the balancing parameter $\alpha$.

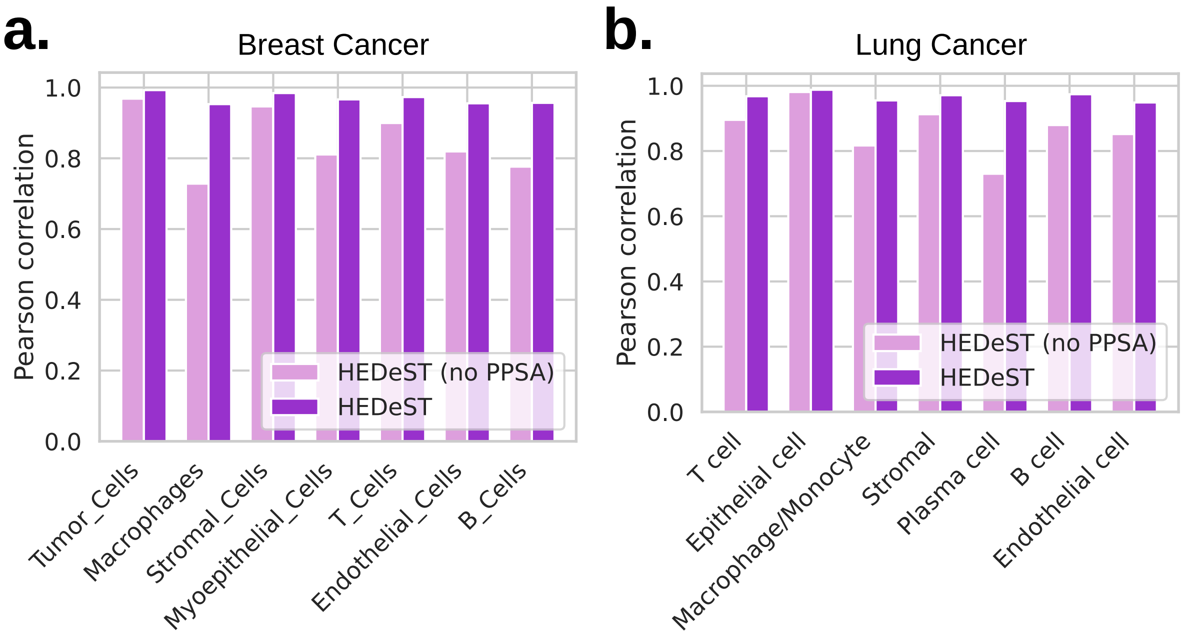

**Fig. 17** Spot-level performance of HEDeST on semi-simulated datasets. HEDeST (no PPSA), HEDeST without PPSA. For each cell type, Pearson correlation between predicted and true proportions was computed before and after PPSA for Breast (**a**) and Lung (**b**), showing that PPSA readjusts probabilities to better match local priors.

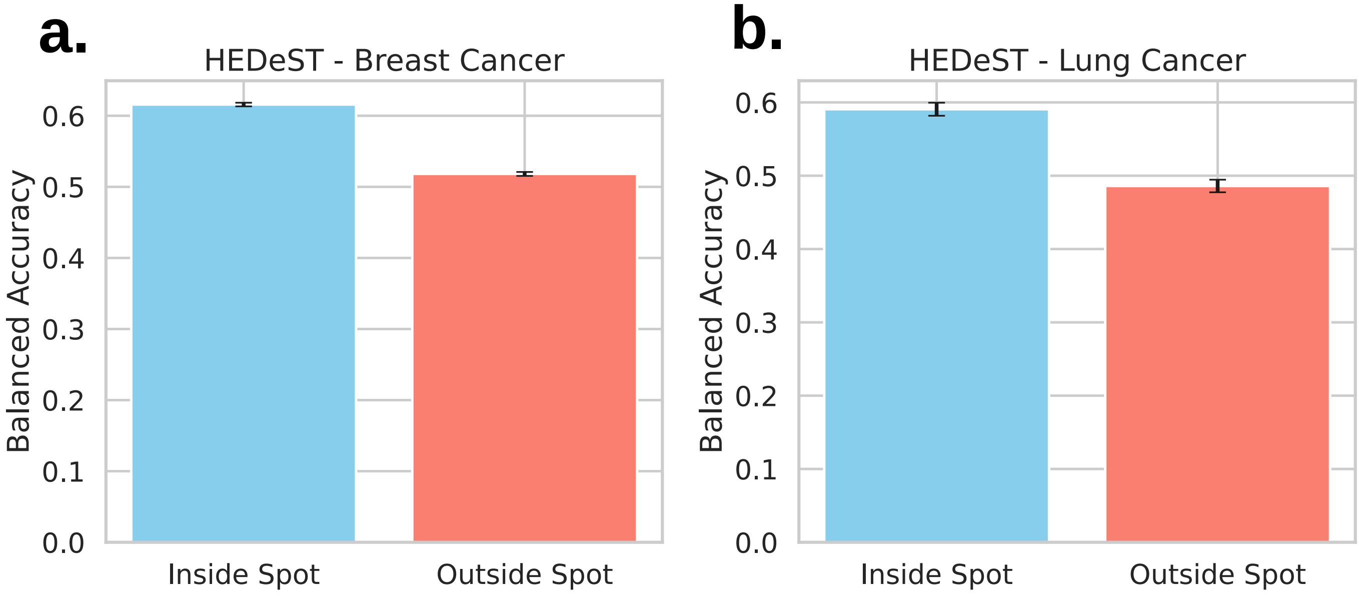

**Fig. 18** Balanced accuracy of cell type predictions inside vs outside spatial spots. **a** Human Breast Cancer dataset. **b** Human Lung Cancer dataset. Bars represent mean balanced accuracy across 10 model seeds, with error bars indicating 95% confidence intervals.

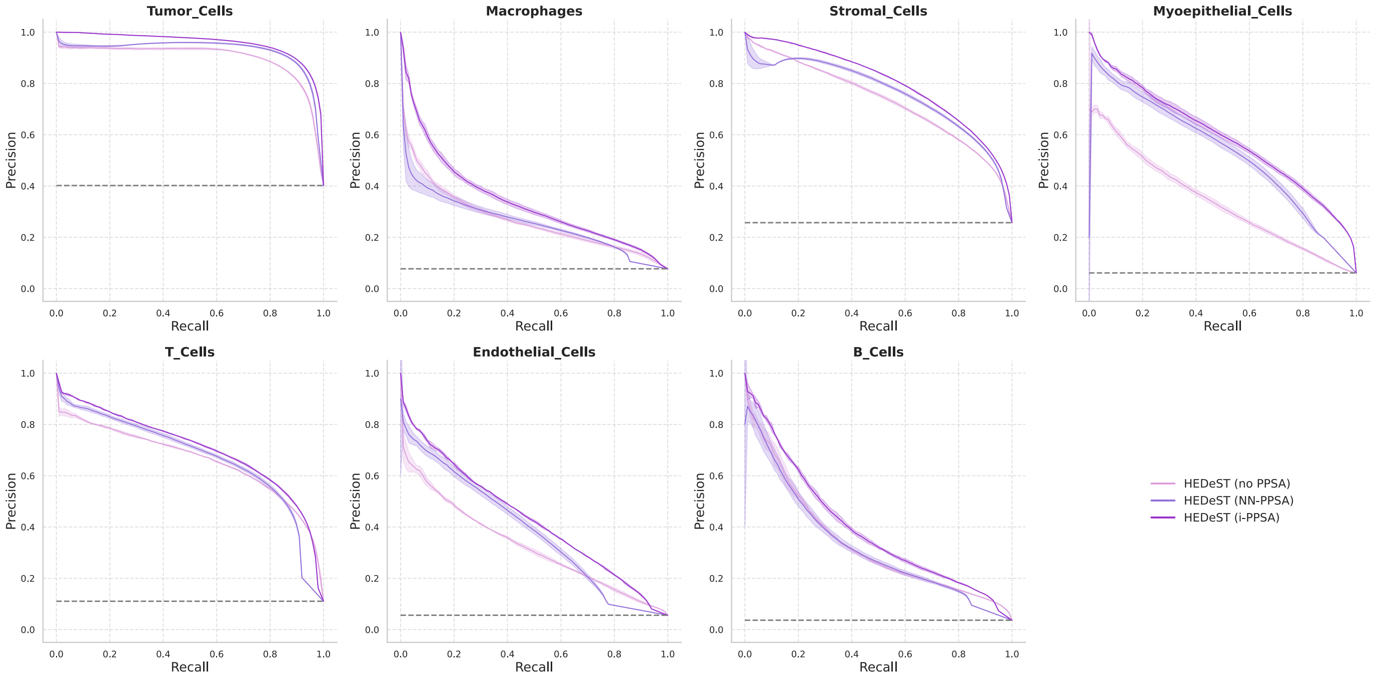

**Fig. 19** Precision-recall curves for cell type predictions on the Xenium Breast Cancer dataset. HEDeST (no PPSA), HEDeST without PPSA. Curves show the trade-off between precision and recall for each cell type across different classification thresholds.

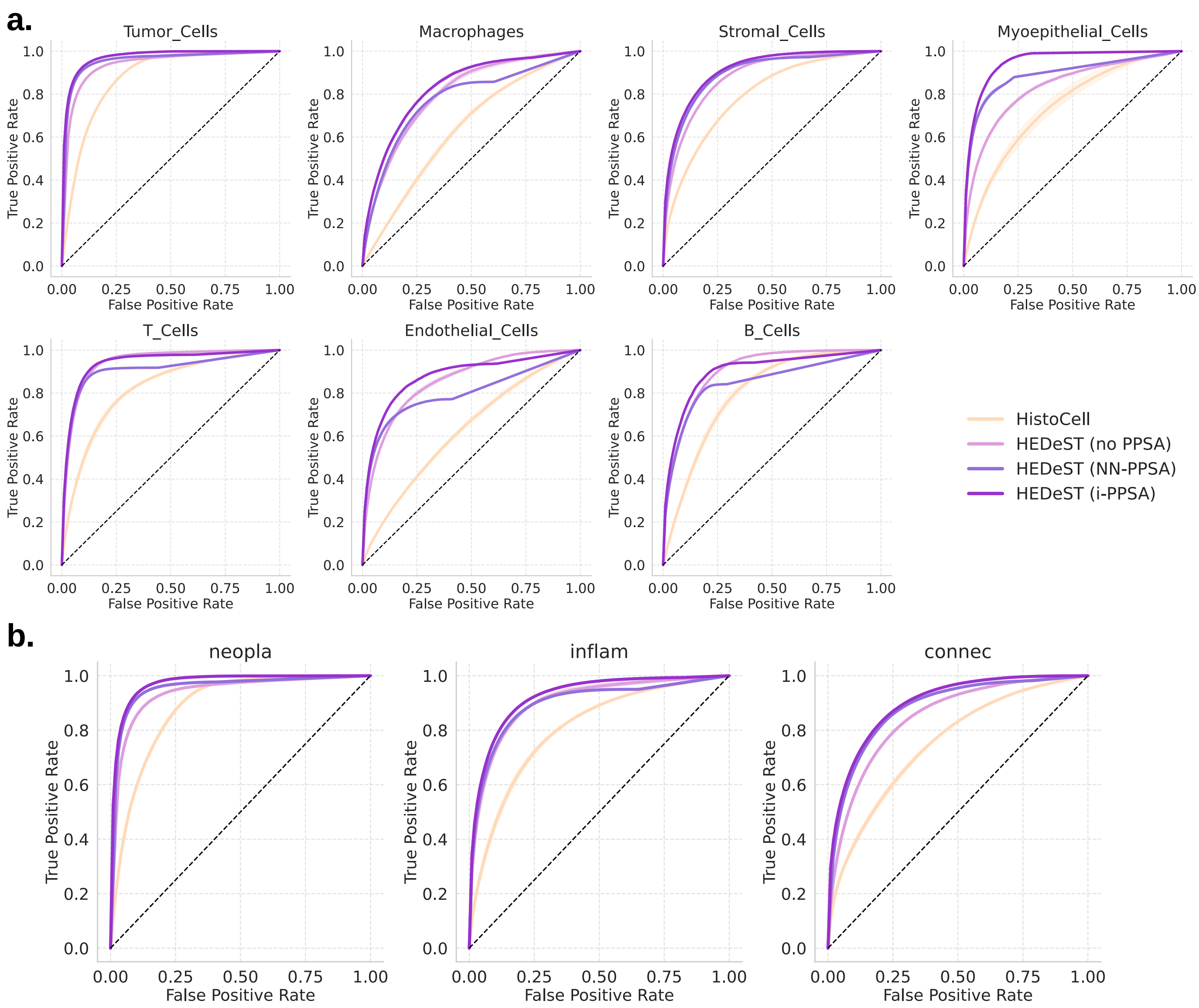

**Fig. 20** ROC curves for cell type predictions on the Xenium Breast Cancer dataset. HEDeST (no PPSA), HEDeST without PPSA; neopla, Neoplastic cells; inflam, Inflammatory cells; connect, Connective cells. **a** ROC curves under standard prediction conditions comparing HEDeST variants with HistoCell. **b** ROC curves restricted to three broad cell type categories.

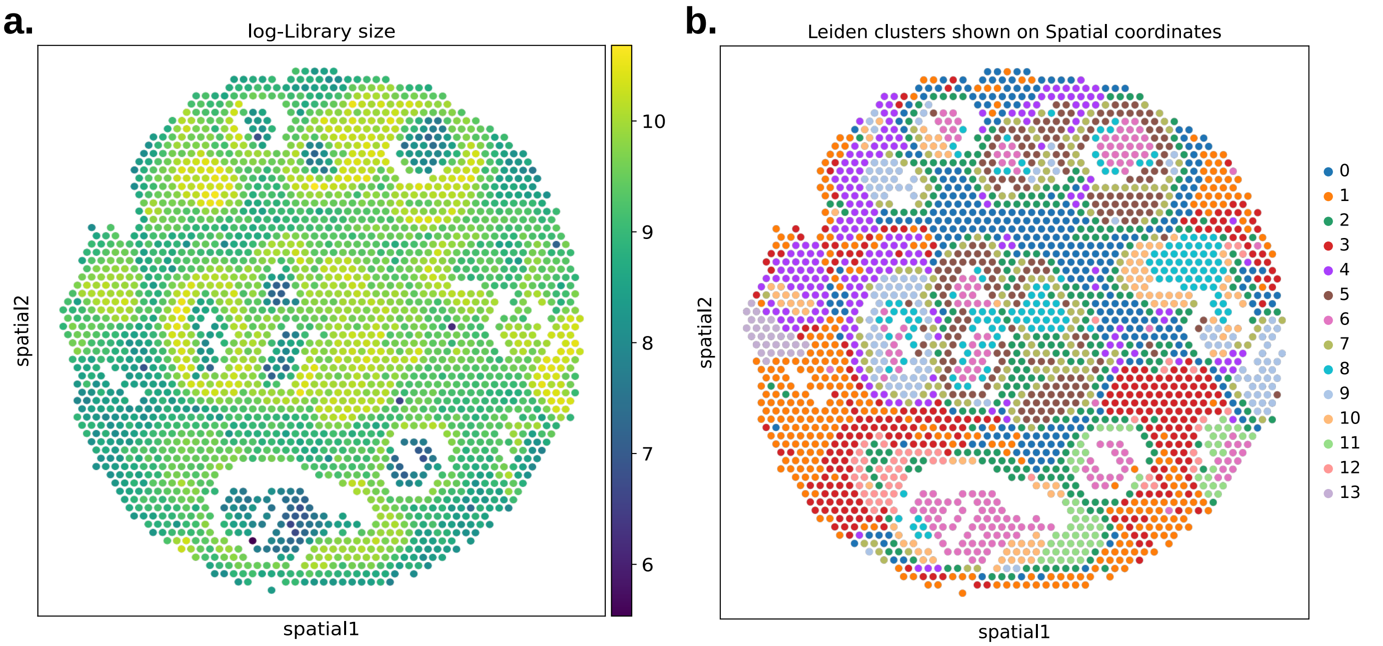

**Fig. 21** Visualization of Visium Breast Cancer data. **a** Spatial spots colored by log-library size, showing variations in total transcript counts across the tissue. **b** Spatial spots colored by Leiden clustering, revealing distinct spatial domains and tissue regions based on transcriptional similarity.

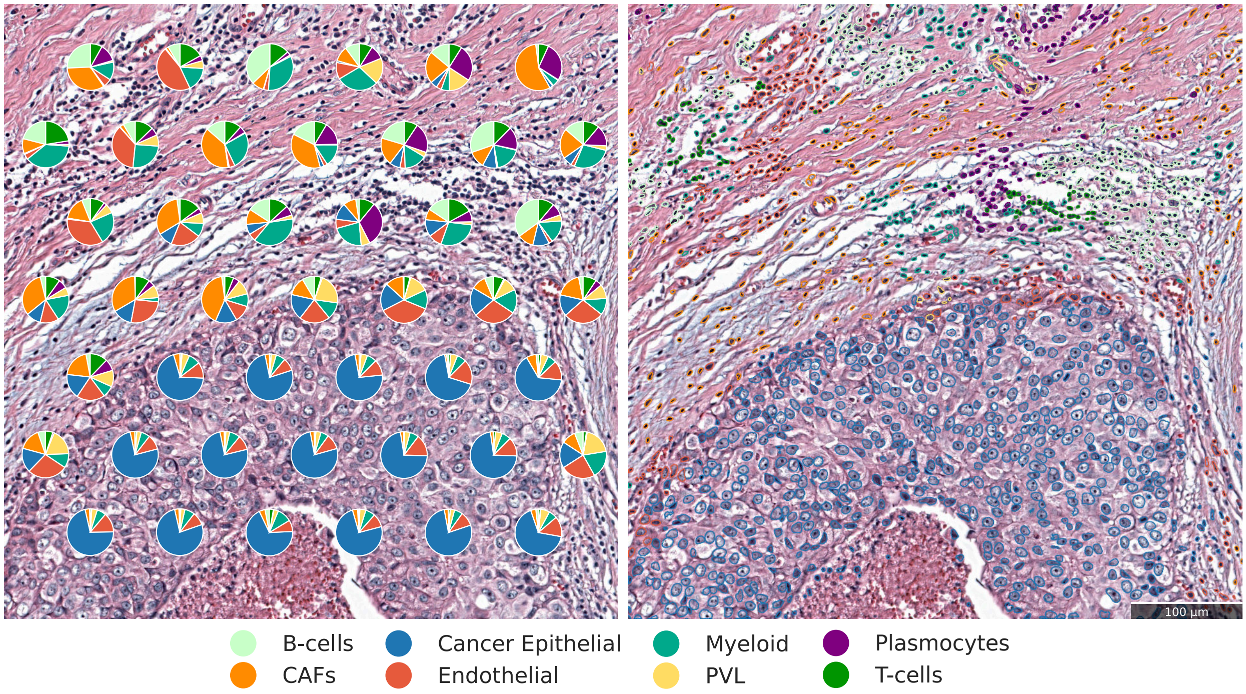

**Fig. 22** Visualization of Visium Breast Cancer slide before and after HEDeST predictions. CAFs, Cancer-Associated Fibroblasts; PVL, Perivascular-like cells. **Left** Cell-type proportions derived from deconvolution, showing spatial distribution of estimated cell-type abundances across tissue spots. **Right** Single-cell annotations displaying cell-type classifications for individual cells within the spatial context.

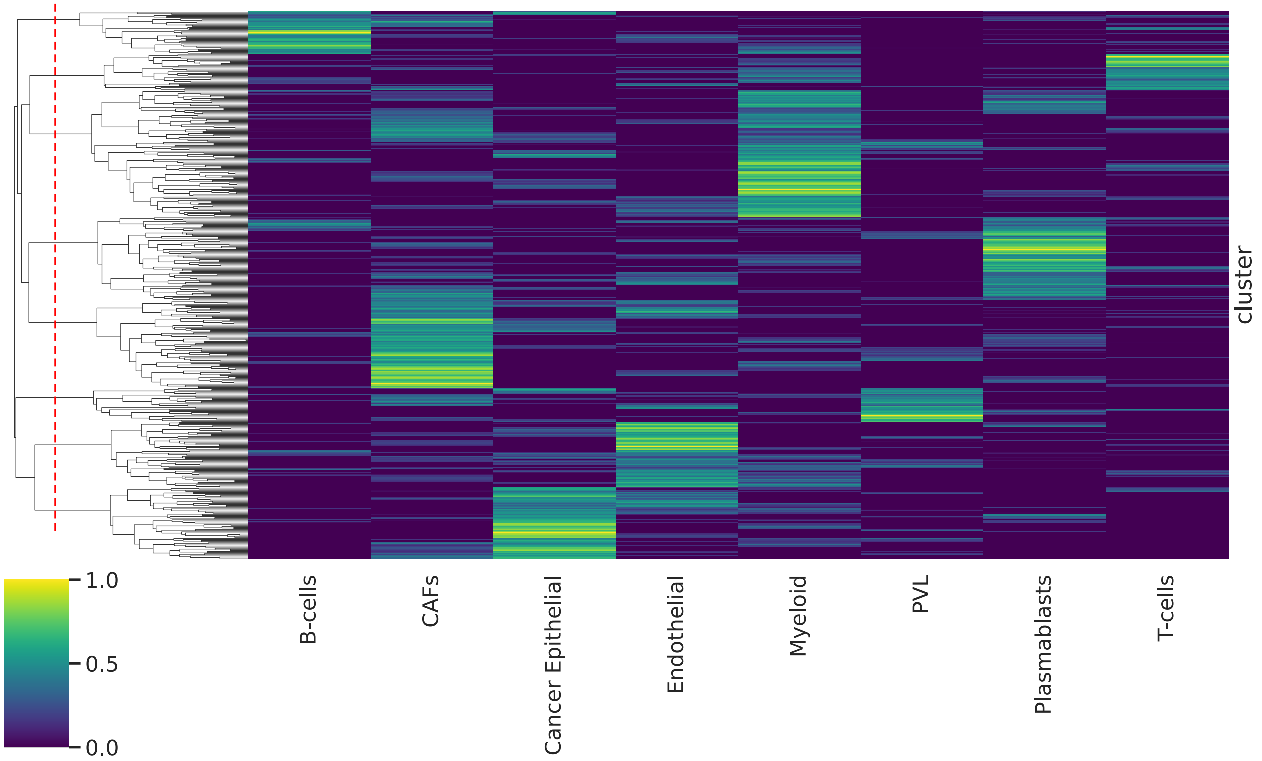

**Fig. 23** Heatmap with dendrogram showing spatial neighborhood analysis of the Visium Breast Cancer dataset. CAFs, Cancer-Associated Fibroblasts; PVL, Perivascular-like cells. Rows represent individual cells, columns represent cell types, and color intensity indicates the proportion of each cell type in the local microenvironment surrounding each cell. The dotted red line indicates the cut height used to define the selected number of clusters.

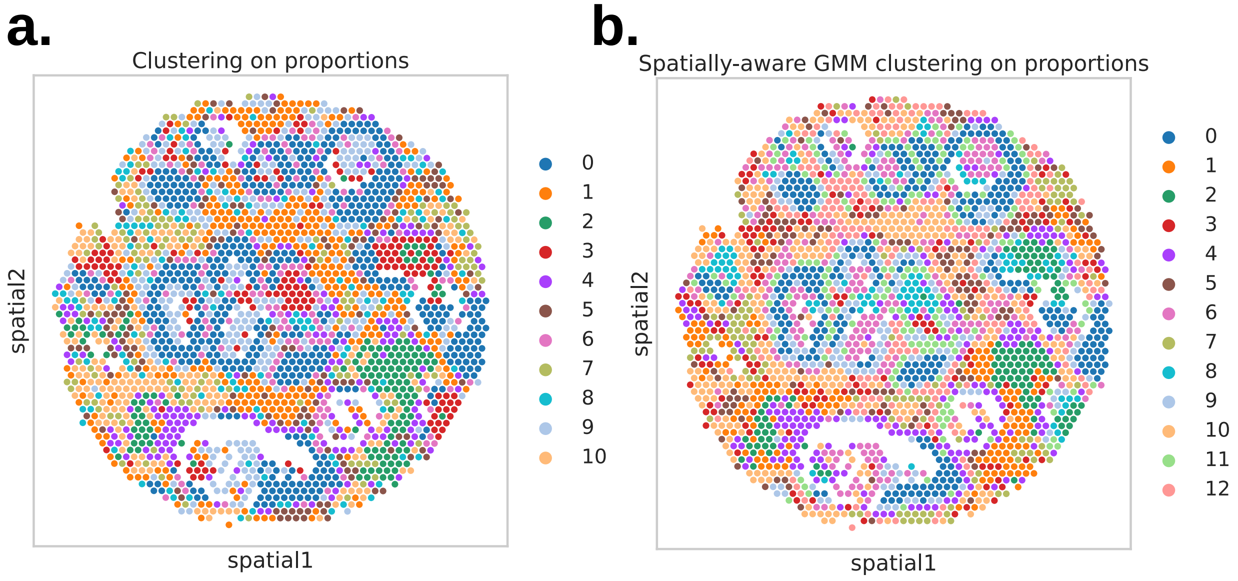

**Fig. 24** Clustering of breast cancer spots based on cell type proportions. **a** Clustering using only spot-level cell type proportions without spatial context. **b** Clustering using spot-level proportions augmented with spatial information from neighboring spots.

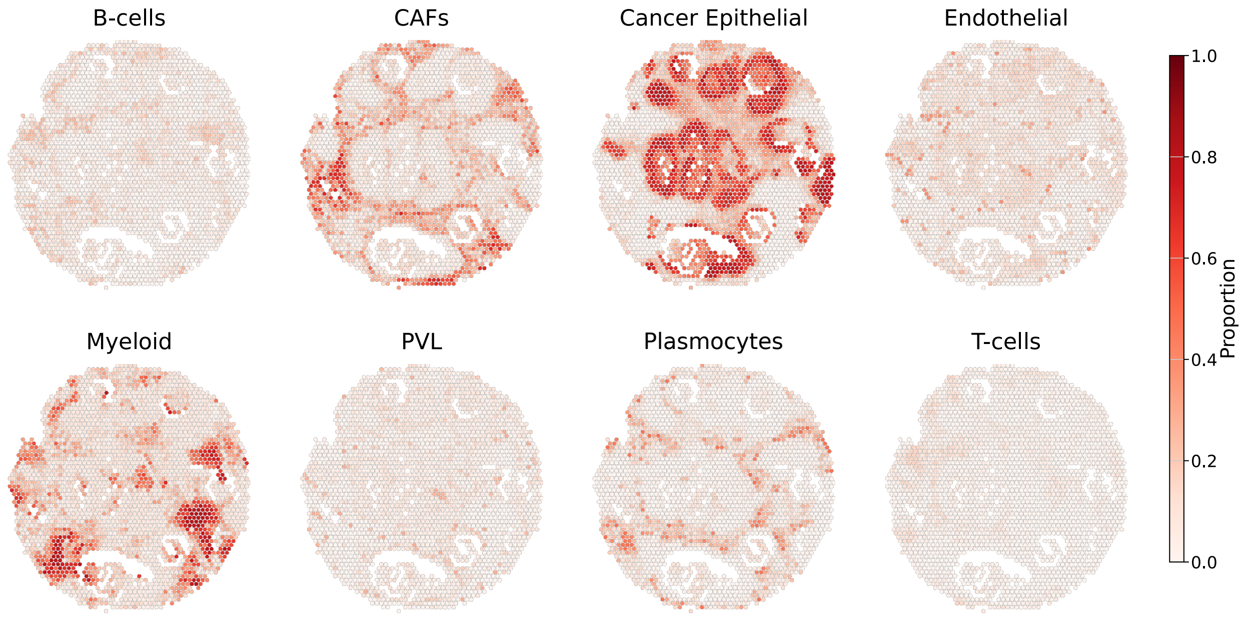

**Fig. 25** Raw cell-type proportions predicted by deconvolution with DestVI [2]. CAFs, Cancer-Associated Fibroblasts; PVL, Perivascular-like cells.

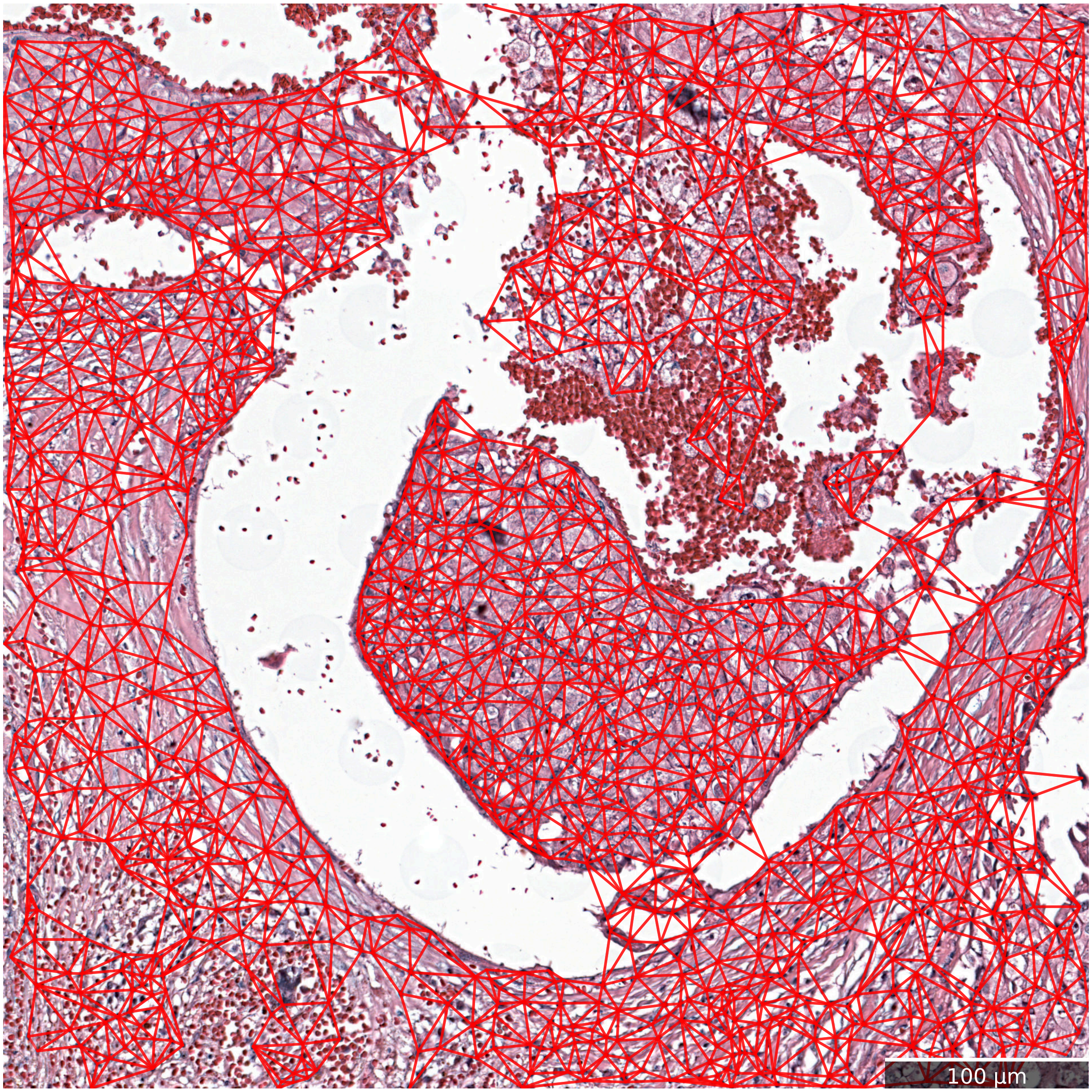

**Fig. 26** Visualization of the Delaunay graph constructed from single-cell coordinates in the Visium Breast Cancer dataset.
